# MicroRNA miR-128 represses LINE-1 (L1) retrotransposition by downregulating the nuclear import factor TNPO1

**DOI:** 10.1101/195206

**Authors:** Adam Idica, Evgueni A Sevrioukov, Dimitri G Zisoulis, Matthias Hamdorf, Iben Daugaard, Pavan Kadandale, Irene Pedersen

## Abstract

Repetitive elements, including LINE-1 (L1), comprise approximately half of the human genome. These elements can potentially destabilize the genome by initiating their own replication and reintegration into new sites (retrotransposition). In somatic cells, transcription of L1 elements are repressed by distinct molecular mechanisms including DNA methylation and histone modifications to repress transcription. Under conditions of hypomethylation (*e.g.* in tumor cells) a window of opportunity for L1 de-repression arises and additional restriction mechanisms become crucial. We recently demonstrated that the microRNA miR-128 represses L1 activity by directly binding to L1 ORF2 RNA. In this study, we tested whether miR-128 can also control L1 activity by repressing cellular proteins important for L1 retrotransposition. We found that miR-128 targets the 3’UTR of the nuclear import factor transportin 1 (TNPO1) mRNA. Manipulation of miR-128 and TNPO1 levels demonstrated that induction or depletion of TNPO1 affects L1 retrotransposition and nuclear import of an L1-RNP complex (using L1-encoded ORF1p as a proxy for L1-RNP complexes). Moreover, TNPO1 overexpression partially reversed the repressive effect of miR-128 on L1 retrotransposition. Our study represents the first description of a protein factor involved in nuclear import of the L1 element and demonstrates that miR-128 controls L1 activity in somatic cells through two independent mechanisms: direct binding to L1 RNA, and regulating a cellular factor necessary for L1 nuclear import and retrotransposition.

## INTRODUCTION

Repetitive elements make up approximately half of the mammalian genomes. A substantial portion of repetitive elements are derived from retrotransposons (long terminal repeats (LTR)-containing and non-LTR), which transpose to new chromosomal locations by reverse transcription of the RNA into DNA, followed by integration of the copied DNA into a new chromosomal location. Retrotransposition of these elements in germ cells lead to integration of new retrotransposons in the genomes of progeny, and since there is no mechanism for excision, they accumulate over evolutionary time scales (1-2).

Long-interspaced nuclear elements-1 (LINE-1 or L1) are the only autonomous transposable elements that are currently active in humans and have directly or indirectly contributed to ∼ 17% of the human genome (1). Intact, active L1 is ∼6 kilobase pairs (kb) in length and contain a 5’UTR, three open-reading frames – ORF1, ORF2 and ORF0 – and a short 3’UTR. The 5’UTR has promoter activity in both the sense and antisense direction (3-6). ORF1 encodes a protein with RNA-binding and nucleic acid chaperone activity and ORF2 encodes a protein with endonuclease and reverse transcriptase activities (2,7-9). ORF0, which is transcribed in the antisense direction, encodes a protein with unknown function, but which enhances L1 activity. L1 mobilizes replicatively from one place in the genome to another by a “copy and paste” mechanism via an RNA intermediate (10-11). L1-RNP complexes have been described to enter the nucleus during cell division (12-13). However, recently L1 retrotransposition has been demonstrated also to take place in non-diving cells such as neurons (14-15). The mechanism by which L1-RNP complexes access the host DNA independently of cell division is unknown.

Integration of retrotransposons at new chromosomal locations can generate new genes and affect expression of already existing genes (16-19). It has been suggested that retrotransposon activity could contribute to various diseases such as neurological disorders and cancer, as well as developmental defects (20-23). As a result, multiple mechanisms have evolved to tightly control retrotransposon activity. In germ cells, specific small RNA subtypes (piRNAs) efficiently counteract L1 activity (24-25). In somatic cells, L1 mobilization is potently inhibited by DNA methylation of the L1 promoter (26-27). However, L1 promoter silencing is greatly attenuated and L1 transcription de-repressed in somatic cells under conditions of hypomethylation, often encountered in cancer cells or in in-vitro reprogramming of somatic cells into induced pluripotent stem cells (iPSCs) (26, 28-29). Under these conditions other mechanisms of L1 restriction are important, including DNA, RNA editing proteins and the microprocessor (AID, APOBECs, ADAR, DGCR8) (30-33).

The recent discovery of microRNAs (miRNAs or miRs) has revolutionized our understanding of gene control. miRs exemplify the emerging view that non-coding RNAs (ncRNAs) may rival proteins in regulatory importance. The majority of the human transcriptome is believed to be under miR regulation, positioning this post-transcriptional control mechanism to regulate many gene pathways (32, 34). miRs function as 21–24-nucleotide (nt) guides that regulate the expression of mRNAs containing complementary sequences. The mature miR is loaded onto specific Argonaute (Ago) proteins, which are then referred to as a miR-inducing silencing complex (miRISC) (34). In animals, partial pairing between a miR and an mRNA target site usually results in reduced protein expression through a variety of mechanisms that involve mRNA degradation and translational repression (35-36). The best-characterized feature determining miR-target recognition are six nucleotide “seed” sites in the 3’UTR of mRNA targets, which perfectly complement the 5’ end of the miR (positions 2-7) (35).

We recently discovered that miR-128 represses activity of L1 retrotransposons in somatic cells, analogous to the role of piRNAs in germ cells. We found a novel mechanism for this regulation in that miR-128 binds directly to L1 RNA in the ORF2 coding region sequence (CRS), resulting in L1 repression (37). In contrast, miRs typically are thought to repress multiple cellular mRNAs by binding to homologous target sequences; the proteins of these target mRNAs often work in concert, so miRs can fine-tune specific cellular networks (38-41).

In this study, we explored if miR-128 also regulates L1 activity in somatic cells by repressing cellular proteins important for its retrotransposition. Here we report that miR-128 significantly represses retrotransposition, by targeting the nuclear import factor Transportin-1 (TNPO-1). TNPO1, also referred to as Karyopherin-ß2 or Importin-ß2, acts by binding to diverse nuclear localization sequences including (PY-NLSs) (37-39). TNPO1-mediated nuclear import requires RanGTP for cargo delivery into the nucleus (46) and known TNPO1 cargoes include viral, ribosomal and histone proteins (45-46). We have determined that miR-128 targets the TNPO1 3’UTR and represses expression of TNPO1 mRNA and protein. In addition, we find that TNPO1 facilitates L1 mobilization and that miR-128-induced TNPO1 deficiency represses L1 retrotransposition, by inhibiting nuclear import of L1-RNP (using ORF1p as a proxy for L1-RNP complexes). This represents the first description of a cellular host factor likely to be involved in nuclear import of L1. Thus, in summary we have discovered a dual mechanism by which miR-128 controls L1 mobilization in somatic cells.

## RESULTS

### miR-128 repress L1 activity

We recently determined that miR-128 directly targets L1 RNA and represses *de novo* retrotransposition and integration in somatic cells including cancer cells, cancer initiating cells (CICs) and induced pluripotent stem cells (iPSCs), which are all characterized by global demethylation and enhanced opportunity for L1 de-repression (37). After demonstrating an important role for miR-128 in the control of L1 retrotransposition in a panel of different cell lines and in iPSCs, we wished to further characterize the mechanism(s) of miR-128-induced restriction of L1 mobilization.

First we initiated analysis to dissect the direct (L1 RNA) versus potential indirect (cellular factors) effects of miR-128 on L1 retrotransposition. We performed colony formation assays using different variants of a neomycin reporter constructs encoding the full length L1 mRNA and a retrotransposition indicator cassette. Briefly, one construct consists of a neomycin gene in the antisense orientation relative to a full-length L1 element, which is disrupted by an intron in the sense orientation (see Supplemental Figure S1A). The neomycin (neo) protein can be translated into a functional enzyme only after L1 transcription and splicing of the mRNA, reverse transcription followed by integration of the spliced variant into the genome, thus allowing the quantification of cells with new retrotransposition events in culture. In addition, we generated a miR-128 resistant variant of the L1 plasmid, by introducing a silent mutation in the miR-128 binding site (in the ORF2 sequence) attenuating miR-128 binding, but allowing L1 to retrotranspose (as described in (37) and see Supplemental Figure S1B). A third variant of the L1 plasmid described by (51) encodes a L1 RNA harboring a D702A mutation in the RT domain of the ORF2 protein, rendering the encoded L1 RT deficient (RT dead). This plasmid variant was used as a negative control (see Supplemental Figure S1C).

miR-128, anti-miR-128 or miR control shRNAs were cloned into the pMIR-ZIP plasmid, packaged into high-titer lentiviruses, and HeLa cells were transduced, puromycin selected and modulation of miR-128 expression levels were verified by miR specific qRT-PCR (Supplemental Figure S2A). miR expressing HeLa cell lines were transfected with either the wildtype (WT), the miR-128 resistant L1 (Mutant) or the RT deficient L1 (RT-dead) neomycin reporter and selected for 14 days with neomycin replenished daily. We verified that the L1 plasmid was introduced into miR-expressing HeLa cells at equal levels by quantifying levels of a neomycin-expressing construct (see Supplemental Figure S2B). We then compared the effect of miR-128 and anti-miR-128 on L1 mobilization to a panel of miR controls (miR-control (miR-control 1), anti-miR-control (miR-control 2) and miR-127, which does not affect L1 retrotransposition (miR-control 3)). In agreement with our previous findings, we observed a significant decrease in the number of neomycin-resistant colonies in cells overexpressing miR-128 and conversely a significant increase in neomycin resistant colonies in anti-miR-128 overexpressing cells (in which endogenously expressed miR-128 is neutralized), relative to HeLa cells with endogenous miR-128 levels, indicating lower versus higher rates of active retrotransposition of WT L1, in cells where miR-128 is either overexpressed or neutralized (Figure 1A left panel)(32). Next analysis of miR-128’s regulation of miR-128-resistant L1 retrotransposition (Mutant) was performed to evaluate potential indirect regulation of L1 by miR-128. We found that miR-128-induction significantly repressed mobilization of miR-128-resistant L1, and that miR-128 neutralization by anti-miR-128, significantly enhanced mobilization of miR-128 resistant L1, relative to miR control (Figure 1A, middle panel). Importantly, miR-modulated HeLa cells encoding RT-deficient L1 (RT-dead) resulted in no neomycin-resistant colonies, demonstrating that colonies obtained upon wild-type and miR-128-resistant L1 plasmid transfections and neo selection are the consequence of a round of *de novo* L1 (Figure 1A, right panel) (48, 51). These results support the idea that miR-128 functions through direct binding of L1 RNA and by regulating cellular co-factors, which L1 is dependent on for successful mobilization.

**Figure 1:**
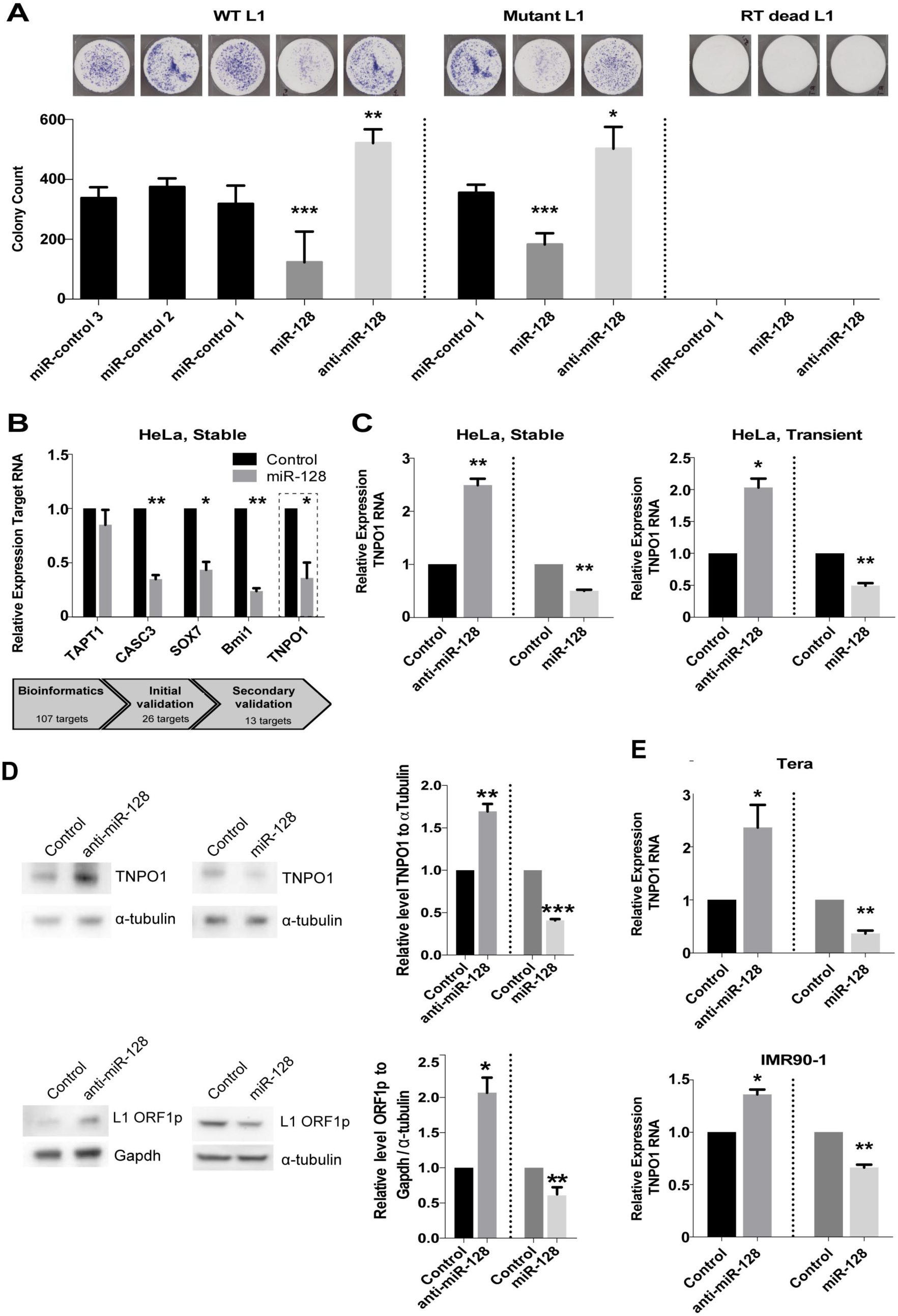
Identification and verification of TNPO1 as a cellular target of miR-128. **(A)** Change in colony count of neomycin-resistant foci was used to determine the level of active retrotransposition in HeLa cells stably transduced with lentiviral constructs encoding a control miRs (control-1, -2, -3), anti-miR-128 or miR-128 transfected with L1 expression plasmid (wild-type L1, left panel). Colony formation assays were performed as described above using a miR-128 resistant L1 expression plasmid (Mutant) or reverse transcriptase incompetent L1 expression plasmid (RT dead L1). Shown as mean ± SEM (n=3, independent biological replicates, *, p<0.05, **, p<0.01). **(B)** Schematic of miR-128 qPCR screen approach. HeLa cells were transiently transfected with miR-128 or control miR mimic, cells were harvested after 72 hrs, RNA was isolated and qPCR was performed for predicted miR-128 targets using GAPDH as housekeeping gene (top panel). Thirteen targets were validated as downregulated in miR-128 treated cells (Supplemental Figure 3) and relative levels of five targets, TAPT1, CASC3, SOX7, Bmi1, and TNPO1 RNA normalized to B2M are shown as mean ± SEM (n=3, IBR, *, p<0.05, **, p<0.01) (bottom panel). **(C)** Relative levels of TNPO1 RNA normalized to B2M in HeLa cells stably transduced or transiently transfected with control miR, anti-miR-128 or miR-128 are shown as mean ± SEM (n=3, independent biological replicates, *, p<0.05, **, p<0.01) **(D)** HeLa cells were stably transduced with control, anti-miR-128 or miR-128 lentiviral constructs and western blot analysis were performed for TNPO1 (top panel), L1 ORF1p (bottom panel), alpha-Tubulin, or GAPDH protein. One representative example of three is shown. Quantification of results (n=3) normalized to Tubulin (TNPO1) or GAPDH (L1 ORF1p) are shown (right panels). **(E)** Relative levels of TNPO1 RNA normalized to B2M were determined in a teratoma cell line (Tera) stably transduced with control miR, anti-miR-128 or miR-128 and iPSCs (IMR90-1) transiently transfected with control miR, anti-miR-128 or miR-128 mimics (n=3, independent biological replicates, *, p<0.05, **, p<0.01). Throughout the figure, *P<0.05; **P<0.01 by two-tailed Student’s *t* test. Uncropped versions of blots are shown in Supplementary Figure 5.

### Identification of miR-128 targets involved in regulation of L1 retrotransposition

miRs often exert their regulatory roles of complex cellular functions by repressing multiple targets in the same signaling pathway. As such miRs can be thought of as master RNA regulators, similar to transcription factors, which are DNA regulators. We have employed different strategies to identify miR-128 targets, which may work in synergy with direct L1 RNA targeting, to limit L1 mobilization. We performed an unbiased screen to validate bioinformatically predicted miR-128 targets by using PicTar and TargetScan (47, 52) (Figure 1B, top panel). HeLa cells were transfected with miR-128 or control miR mimics, 107 targets were analyzed by qRT-PCR and 13 potential miR-128 targets were verified twice, including TAPT1, CASC3, SOX7, BMI-1 and TNPO1 (Figure 1B, bottom panel and Supplemental Figure S3).

An area of L1 biology which the literature is conflicted about, deals with whether L1 ribonuclear (L1-RNP) complexes are dependent on cell division for nuclear import or not (12-13,15). Interestingly, Macia et al. recently demonstrated that L1 can retrotranspose efficiently in mature nondividing neuronal cells, however the mechanism responsible for active nuclear import is unknown (14). With this in mind, we were excited to identify Transportin-1 (TNPO1) as a potential miR-128 target, as TNPO1 functions in nuclear import of a variety of RNA binding proteins critical for various steps in gene expression (56, 59,).

We determined that stably transduced HeLa cells expressing anti-miR-128 exhibit significantly higher levels of TNPO1 mRNA relative to the control sequence (Figure 1C, left panel), in contrast to miR-128 overexpressing HeLa cells in which TNPO1 mRNA was significantly reduced (Figure 1C, left panel). To rule out the possibility that the observed miR-128 effect was an artifact stemming from genomic integration of lentiviral encoded miRs, we transiently transfected miR-128, anti-miR-128 or control miR mimic oligonucleotides into HeLa cells, as an alternative approach and verified the effect of miR-128 and anti-miR-128, relative to miR controls (Figure 1C, right panel and Figure 1B, bottom panel). Next we determined that miR-128 versus anti-miR-128 regulated the protein level of TNPO1 correlating with the observed changes in expression levels of TNPO1 mRNA (Figure 1D, top panel, quantifications top right panel) and that these changes were accompanied by significant ORF1p reductions versus increases (Figure 1D, bottom panel, quantification bottom right panels and (37)). Finally, to exclude the possibility that miR-128 exclusively targets TNPO1 mRNA in HeLa cells we tested a teratoma cell line (Tera-1) and an induced pluripotent stem (iPS) cell line (IMR90). We found that TNPO1 mRNA expression levels were significantly changed in Tera-1 and IMR90 cells, in addition to HeLa cells (Figure 1E and Figure 1C). These combined results show that miR-128 regulates the expression levels of TNPO1 in different cell types.

### miR-128 interacts with a target sequence in the 3’UTR of TNPO1 mRNA

Next we wished to examine whether miR-128 indirectly regulates TNPO1 expression or directly interacts with TNPO1 mRNA. Bioinformatics analyses identified 3 potential seed matches in TNPO1 mRNA (Figure 2A). TNPO1 3’UTR or coding reading frame sequence including the three potential miR-128 binding sites, were cloned into a luciferase-based miR-binding site reporter construct. In addition, a perfect 23 nt miR-128 sequence (positive control) luciferase construct was generated. HeLa cells were transfected with one of the TNPO1 binding site-encoding plasmids in addition to mature miR-128 or miR control mimics. Luciferase activity was significantly reduced in cells transfected with miR-128 and encoding binding site #1 (8-mer perfect seed site in 3’UTR) (See Figure 2A and Figure 2B). In contrast miR-128 expression did not substantially reduce luciferase activity in cells encoding binding site #2 or #3. These results indicate that miR-128 preferentially targets TNPO1 mRNA by binding to site #1.

**Figure 2:**
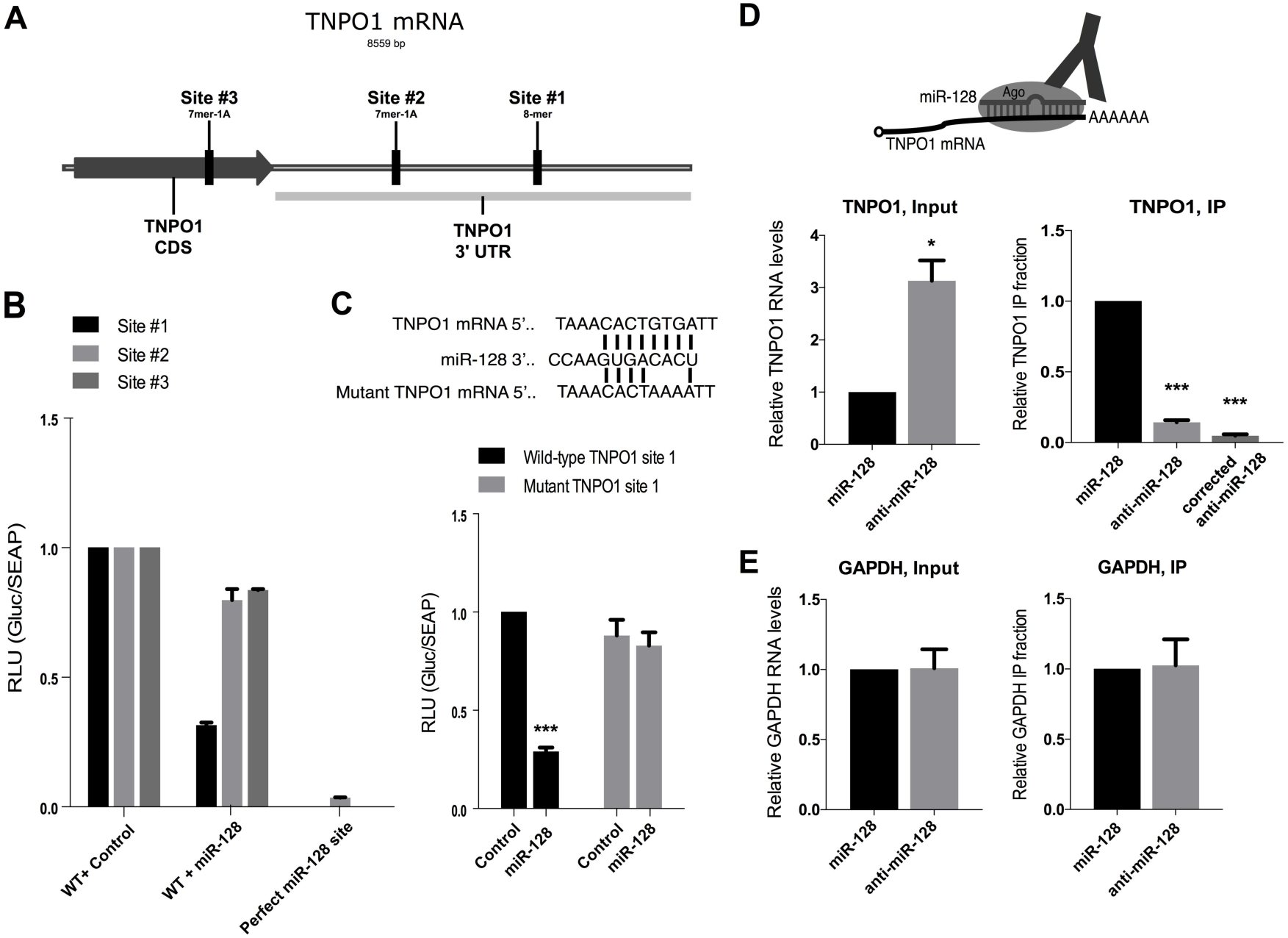
miR-128 represses TNPO1 by binding directly to the 3’UTR of TNPO1 mRNA. **(A)** Schematic of the three predicted miR-128 binding sites in the TNPO1 mRNA (coding region CDS and 3´UTR are shown). miR-128 binding site #1 in TNPO1 3’ UTR is a perfect 8-mer seed site, site #2 (in 3’UTR) and site #3 (in CRS) are both 7-mer seed binding sites **(B)** Relative luciferase levels of HeLa cells transfected with constructs expressing a luciferase gene fused to the wild-type (WT) binding sequence for site 1, 2, 3, or positive control sequence corresponding to the 22 nucleotide perfect match of miR-128 along with transfections of control or miR-128 mimics were determined 48hr post-transfection. Results shown as mean ± SEM (n=3, independent biological replicates, *, p<0.05, ***, p<0.001). **(C)** Schematic of miR-128 binding to WT TNPO1 3’UTR mRNA or mutant seed site TNPO1 mRNA (top panel). Relative luciferase levels of HeLa cells transfected with the reporter plasmid for WT site 1 or mutated site 1, co-transfected with control miR or miR-128 mimics were determined 48hr post-transfection. Results shown as mean ± SEM (n=3, independent biological replicates, ***, p<0.001). **(D)** Schematic representation of the Ago immunopurification strategy of miR-128-TNPO1 mRNA complexes (Ago-RIP) (top panel). HeLa cell lines are generated where miR-128 is either stably neutralized (by anti-miR-128) or over-expressed. Relative expression of TNPO1 mRNA normalized to B2M is shown for input samples (left panel); relative fraction of TNPO1 transcript levels associated with Ago complexes is shown for IP samples (right panel). TNPO1 IP fractions normalized to the levels of TNPO1 in input are shown as “corrected” (right panel). Results shown as mean ± SEM (n=3, independent biological replicates, *, p<0.05, ***, p<0.001). **(E)** Relative levels of GAPDH in the same input and IP samples were determined as a negative control. Results are shown as a mean ± SEM. Results shown as mean ± SEM (n=3, independent biological replicates).

Next, mutations were introduced into the putative miR-128 binding site in the TNPO1 mRNA encoding site #1 in 3’UTR (Figure 2A), to determine if this sequence is responsible for the interaction with miR-128 (Figure 2C, top panel). Luciferase activity was again significantly lower than controls in HeLa cells transfected with the wild-type (WT) TNPO1 site #1 plasmid and mature miR-128 supporting the conclusion that miR-128 can bind to the WT TNPO1 3’UTR sequence and prevented the translation of luciferase (Figure 2C, bottom left panel). In contrast, HeLa cells transfected with the mutant TNPO1 3’UTR mutant binding site and mature miR-128 or control miRs exhibited luciferase activity at the same levels as the WT TNPO1 and miR-control cells; consistent with the conclusion that miR-128 could no longer bind and repress reporter gene expression (Figure 2C, bottom right panel).

Furthermore, Argonaute (Ago) complexes containing miRs and target mRNAs were isolated by immuno-purification and assessed for relative complex occupancy by the TNPO1 mRNA to validate that miR-128 directly targets TNPO1 mRNA in cells (Figure 2D, top) in miR-128-versus anti-miR-128-overexpressing HeLa cells, as previously described (37). The relative level of TNPO1 mRNA was significantly lower in cells stably overexpressing miR-128 when compared to those expressing anti-miR-128 constructs, as expected (Figure 2D middle, Input). Despite the increased levels of TNPO1 mRNA (because of lower miR-128 expression levels), which may underestimate the scale of the effect, the relative fraction of Ago-bound TNPO1 mRNA significantly increased when miR-128 was overexpressed (Figure 2D, IP). When correcting for the lower expression level of TNPO1 mRNA, the increase in miR-128 bound TNPO1 mRNA was even more significant (Figure 2D, top right panel). In contrast, miR-128 did not repress GAPDH mRNA expression levels or immuno-purified gapdh mRNA, as expected (Figure 2E). We interpret this to mean that high levels of miR-128 lead to higher levels of TNPO1 mRNA being bound and regulated directly by miR-128. These data support the conclusion that miR-128 represses TNPO1 expression via a direct interaction with the target site located in the 3’UTR of the TNPO1 mRNA.

### TNPO1 modulation regulates L1 activity and de novo retrotransposition

TNPO1 functions by interacting with nuclear localization sequences on protein cargoes and facilitates nuclear import (55-58). We hypothesized that L1-RNP may utilize TNPO1-dependent active transport, in addition to accessing the host DNA during cell division.

First we wished to evaluate if TNPO1 directly plays a role in L1 mobilization. For this purpose we generated TNPO1 constructs expressing TNPO1 shRNA (to obtain TNPO1 knock-down HeLa cells), or encoding the full-length TNPO1 mRNA transcript harboring the 5.6 kb 3’UTR including the miR-128 binding site (to generate HeLa cells overexpressing TNPO1) and control plasmids. We verified that TNPO1 shRNA or overexpression plasmids significantly reduced versus increased mRNA of TNPO1, relative to controls (Figure 3A). We also evaluated whether TNPO1 knockdown or overexpression are toxic to cells or affect cell proliferation. Morphological and cell proliferation analysis of TNPO1 modulated HeLa cells showed that TNPO1 knockdown or overexpression, is not toxic to HeLa cells, which proliferate at a similar rate relative to plasmid control HeLa cells (Supplemental Figure S4A). Since HeLa cells express low levels of endogenous L1 activity, we transiently transfected a construct encoding the full-length wild-type (WT) L1 and monitored the effects of TNPO1 depletion on artificially expressed L1 mRNA. We then performed colony formation assays to determine a possible requirement of TNPO1 on new L1 retrotransposition events. We verified that the L1 plasmid was introduced into TNPO1-expressing HeLa cells at similar levels by quantifying levels of neo-encoding expression plasmid (Supplemental Figure S2C and S2D). We then determined that cells deficient in TNPO1 exhibited a significantly lower number of neomycin-resistant colonies, versus cells overexpressing TNPO1, which showed a significant increase in neomycin-resistant colonies relative to controls (Figure 3B). This is consistent with lower versus higher rates of *de novo* retrotransposition and genomic integration (Figure 3B, shown as colony counts (%) and colony counts). TNPO1-modulated HeLa cells encoding RT-deficient L1 (RT-dead) resulted in no neomycin-resistant colonies, demonstrating that colonies obtained upon wild-type L1 plasmid transfections and neo selection are the consequence of a round of *de novo* L1 (data not shown). Next, protein lysates from TNPO1 deficient cells and TNPO1 overexpressing cells were prepared and TNPO1 and ORF1p protein levels were found to be significantly reduced in TNPO1 deficient cells, and increased in TNPO1-induced cells, as compared to controls (3C, left panel, quantification, right panel). The amount of L1 mRNA (ORF2) was regulated by TNPO1 consistent with the observed effect on L1 protein (ORF1p) abundance (data not shown). We noticed that the global amount of L1 protein changes when TNPO1 levels changes. This may be a consequence of accelerated degradations of L1-RNP components, caused by dysregulated nuclear transport of L1.

**Figure 3:**
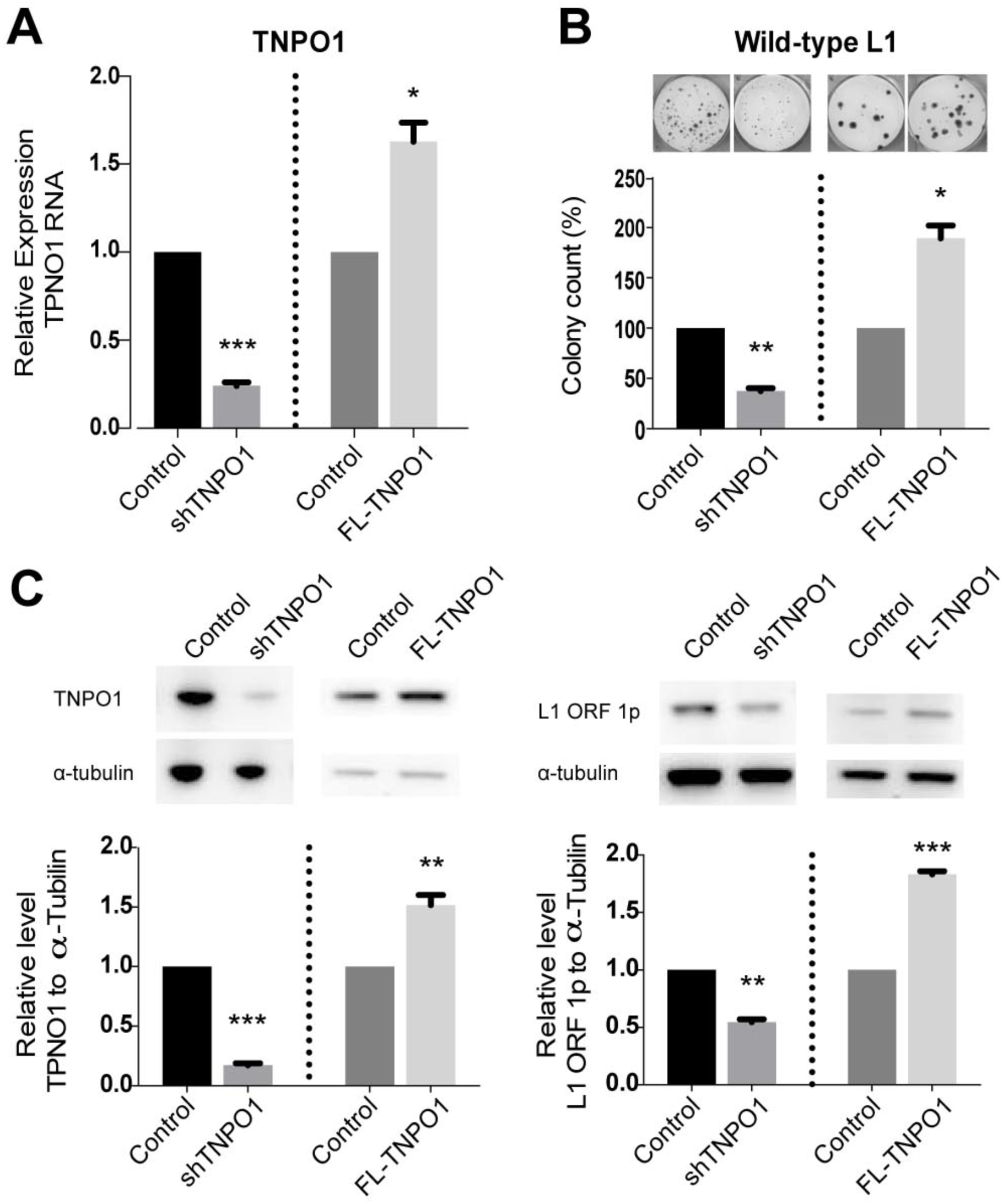
TNPO1 knock-down reduces L1 activity, whereas TNPO1 overexpression enhances L1 retrotransposition. **(A)** Relative expression of TNPO1 RNA normalized to B2M in the same samples was determined (right panel). Results shown as a mean ± SEM (n=3, independent biological replicates, *, p<0.05, ***, p<0.001). **(B)** *de novo* retrotransposition was determined by quantification of neomycin-resistant foci of HeLa cells stably transfected with plasmids encoding Controls (control), shTNPO1, FL-Control or FL-TNPO1 co-transfected with L1 expression plasmid (Wild-type L1). Shown as mean ± SEM (n=3, independent biological replicates, *, p<0.05, **, p<0.01). **(C)** Relative expression of amount of ORF2 normalized to B2M in HeLa cells stably transfected with a shControl (Control), shTNPO1, FL-Control (Control), or full-length TNPO1 overexpression (FL-TNPO1) plasmid (left panel). **(D)** Western blot analysis of TNPO1 and alpha-tubulin (α-tubulin) (protein levels in HeLa cells stably transduced with Controls, shTNPO1, or FL-TNPO1 plasmid (left panels). One of three representative examples is shown. Quantification of results (n=3) normalized to alpha-tubulin is shown (right panels). Uncropped versions of blots are shown in Supplementary Figure 5.

These combined data support the conclusion that TNPO1 neutralization or overexpression results in a corresponding decrease or increase in new retrotransposition events and establish a role of TNPO1 as a novel and specific modulator of L1 activity.

### TNPO1 depletion inhibits L1 nuclear import

TNPO1 belongs to the family of transportins, which also includes TNPO2 and TNPO3 (59). All three protein subtypes are expressed in all examined tissues and function in nuclear import (55-59). Reminiscent of the generally accepted role which TNPO3 plays in nuclear import of the pre-integration complex (PIC) of HIV-1 (60-66), we next decided to explore whether TNPO1 functions in a similar manner by assisting in the nuclear import of the L1-RNP complex. Faced with the difficulties of investigating RNA and proteins encoded by endogenous L1s, we developed a construct expressing a tagged protein of L1 containing HA (ORF1p-HA) and used localization of ORF1p as a proxy to reflects localization of L1-RNP, keeping in mind the limitations of this approach. We generated stable TNPO1 overexpressing and TNPO1 knock-down HeLa cell lines, which were transiently co-transfected with full-length wild-type (WT) L1 and ORF1p-HA or control vector and ORF1p localization was visualized and quantified by immunofluorescence confocal analysis. As the FL-TNPO1 plasmid co-expresses GFP, an alternate secondary antibody was used to visualize ORF1p in TNPO1-induced HeLa cell lines (AlexaFluor 568). We determined that TNPO1 reduction (shTNPO1) resulted a significant reduction of nuclear ORF1p (as determined by ORF1p expression in the nucleus as a measure of total cellular ORF1p) (Figure 4A, top panels, quantification top right. White arrows indicate examples of ORF1p nuclear staining in the single channel images) and overexpression of TNPO1 (FL-TNPO1) resulted in a significant increase in localization of ORF1p in the nucleus, as compared to control cells (Figure 4A, bottom panel. Quantification bottom right panel). Untransfected cells are shown in (Supplemental Figure S4C). As a positive control, a known TNPO1 interaction partner, TBP associated factor 15 (TAF15), was also analyzed (67). As expected, TNPO1 knock-down decreased the nuclear localization of TAF15, which was, instead, found in the cytoplasm and at the plasma membrane (see Supplemental Figure S4D).

**Figure 4:**
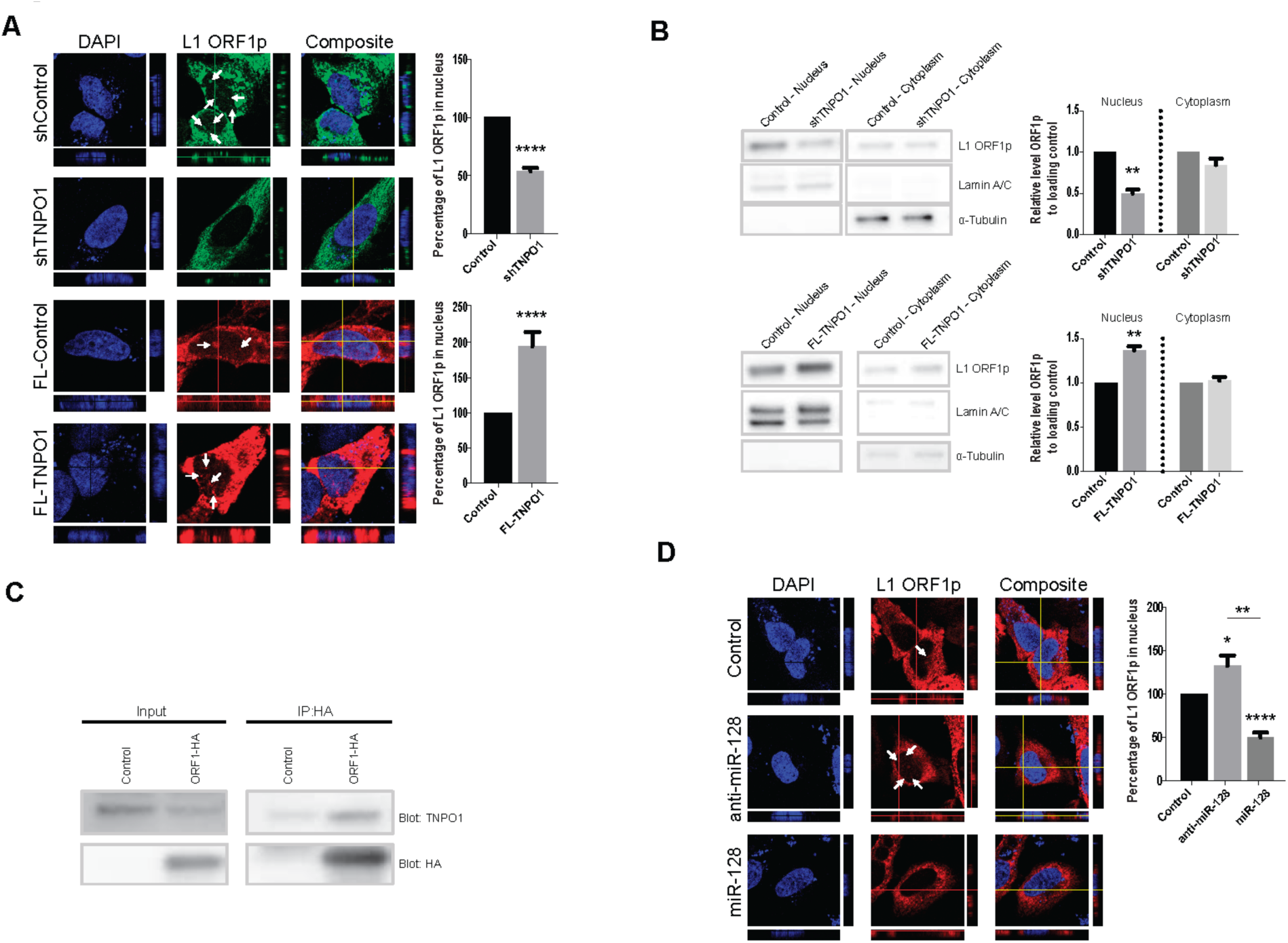
TNPO1 knockdown reduces nuclear import of L1 (ORF1p), whereas induced expression of TNPO1 enhances nuclear import of L1 (ORF1p). **(A)** Localization of L1 ORF1p-HA was determined in HeLa cells stably expressing Controls, shTNPO1 or FL-TNPO1, then co-transfected with full-length wild-type (WT) L1 and ORF1p-HA or control vector. Representative orthogonal views of z-stack images are shown. Quantification of L1 ORF1p-HA localization to the nucleus is shown (represented as a percentage of L1 ORF1p in nucleus/all L1 ORF1p in the image, white arrows indicate examples of ORF1p nuclear staining in the single channel images). Results are shown as the mean percentage of L1 ORF1p in the nucleus ± SEM (n=50, technical replicates of 3 independent biological replicates, ****, p<0.0001). TAF15, a verified TNPO1 cargo, was used as a positive control (see Supplemental Figure 4D). **(B)** Subcellular fractionation analysis was performed on TNPO1 modulated HeLa cells, which was co-transfected with full-length wild-type (WT) L1 and ORF1p-HA or control vector. Western blot analysis of L1-ORF1p-HA, Lamin A/C, or alpha-tubulin (α-tubulin) protein levels in nuclear (N) or cytoplasmic (C) fractions of HeLa cell protein-containing lysates stably expressing Controls, shTNPO1 or FL-TNPO1 (one representative of 3 is shown). Quantification of results (n=3) normalized to Lamin A/C (nuclear), or α-tubulin (cytoplasmic) is shown. **(C)** HeLa cells were transfected with ORF1-HA expression plasmid, HA was immunoprecipitated and co-immunoprecipitated TNPO1 was determine by blotting for native TNPO1. (One representative example out of two is shown). **(D)** Localization of L1 ORF1p-HA was determined in HeLa cells stably transduced with miR control (control), anti-miR-128, or miR-128, then transfected with the ORF1p-HA expression plasmid. Representative orthogonal views of z-stack images are shown. Quantification of L1 ORF1p-HA localization to the nucleus (represented as a percentage of L1 ORF1p in nucleus/all L1 ORF1p in the image, white arrows indicate examples of ORF1p nuclear staining) is shown as the mean percentage of L1 ORF1p in the nucleus ± SEM (n=50, technical replicates of 3 independent biological replicates, *, p<0.05, ****, p<0.0001). Uncropped versions of blots are shown in Supplementary Figure 5.

Next we subjected TNPO1 modulated HeLa cell lines, which were transiently co-transfected with full-length wild-type (WT) L1 and ORF1p-HA, to subcellular fractionation analysis, keeping in mind the limitation of this approach. Nuclear and cytoplasmic fractionations were evaluated by determining the expression levels of α-tubulin (cytoplasmic) and Lamin A/C (nuclear) (Figure 4B, Western blots) and TNPO1 knock-down and overexpression were verified by qRT-PCR (Supplemental Figure S4B). We next examined the effect of TNPO1 modulation on encoded L1 protein (ORF1p) as an indirect measure of L1-RNP localization. Analyzing the ORF1p levels in nuclear versus cytoplasmic fractions from TNPO1 knock-down HeLa cells lines (shTNPO1) (shown in Figure 3C, left panel), showed a substantial decrease in ORF1p levels in the nucleus, relative to controls (Figure 4A, top panels). Overexpression of TNPO1 (FL-TNPO1) resulted in an increased nuclear L1 ORF1p expression as compared to controls (Figure 4B, bottom panels). We did not observe a significant change in ORF1p levels in the cytoplasmic fractions. This is not too surprising as the vast majority of ORF1p is localized in the cytoplasm, thus changes in expression levels might not be measurable, as opposed to expression levels in the nucleus. We noted that ORF1p levels as determined by western blot analysis following subcellular fractionation surprisingly showed a ratio of less ORF1p in the cytoplasm versus nucleus. This finding was in contrast to our confocal analysis of ORF1p localization. This difference is possibly due to a much more dilute cytoplasmic fraction, as compared to nucleic fraction. However, even with these limitations in mind, the combined results from the confocal and subcellular fractionation analysis indicate that TNPO1 is facilitating L1’s access to host DNA.

Furthermore, we performed immunoprecipitation analysis to evaluate if TNPO1 interacts with ORF1p. HeLa cells were transduced with a tagged version of ORF1 (ORF1p-HA), protein-containing lysates were prepared and ORF1p immunoprecipitations were performed and blotted for TNPO1 and HA. The ORF1p co-immunoprecipitations results suggest that ORF1p (L1-RNP complex) and TNPO1 interact (Figure 4C). However, further studies are needed to determine whether this interaction is direct or indirect through an RNA bridge, as previously demonstrated for many ORF1p binding partners (68).

Finally, we evaluated the effect of *miR-128-induced TNPO1 repression* on L1 nuclear import. miR-128, anti-miR-128 or control miR HeLa cells were transfected with ORF1p-HA expression plasmids and localization of L1 was analyzed by confocal analysis, as described above. miR-128-mediated TNPO1 repression (verified and shown in Figure 1D, top panel) resulted in a significant decrease in L1 ORF1p nuclear localization (Figure 4D, right panel. Quantification far right panel), whereas anti-miR-128-induced TNPO1 expression significantly increased nuclear L1 ORF1p expression levels (Figure 4D, middle panel. White arrows indicate examples of ORF1p nuclear staining in the single channel images. Quantification, far right panel).

This body of work supports the idea that miR-128-induced TNPO1 repression results in a modest, but significant and reproducible decrease in nuclear import of some L1-RNP complexes or components of L1-RNP complexes (using ORF1p as a proxy), accumulation of L1 (ORF1p) in the cytoplasm and a significant reduction in L1 retrotransposition events. Additional studies are needed to determine whether functional L1-RNP complexes are actively transported into the nucleus and whether this event is facilitated by TNPO1. In summary, our findings support the idea that in addition to direct access of L1-RNP to host DNA during cell division, some L1-RNP complexes are imported into the nucleus via TNPO1.

### TNPO1 is a functional target of miR-128-induced L1 repression

We have previously demonstrated that miR-128 targets L1 RNA and represses L1 activity by a direct interaction, similar to how miRs represses replication of RNA virus (69, 70). In addition, we have now determined that miR-128 is capable of repressing miR-128 resistant L1 (using a L1 mutant vector), by an indirect mechanism (Figure 1A). With this in mind we wished to evaluate the significance of TNPO1 as a functional mediator of miR-128-induced L1 repression.

We utilized the L1 mutant vector, in which the miR-128 binding site had been mutated and miR-128 is no longer able to bind (miR-128 resistant L1). In addition, in order to perform TNPO1 rescue experiments, we needed to overexpress a miR-128-resistant version of the TNPO1 vector, as miR-128 may otherwise be able to bind to WT TNPO1 plasmid and could in theory function as a miR-128 sponge. We generated a miR-128 resistant full-length TNPO1 vector, in which the miR-128 binding Site#1 in the 3’UTR had been mutated according to our mutation analysis and miR-128 was no longer able to bind (Figure 2C) (FL-TNPO1mut).

We found that overexpression of TNPO1 (WT and miR-128 resistant) in miR-128 overexpressing HeLa cells were able to partially but significantly, rescue miR-128-induced repression of L1 retrotransposition and genomic integration as determined by colony formation assays, relative to controls for WT L1 (Figure 5A, left panel and Supplemental Figure S6A). Similar results were obtained when rescuing miR-128 L1 restriction with TNPO1 (WT and miR-128 resistant) of the Mutant L1 plasmid, relative to controls (Figure 5A, right panel and Supplemental Figure S6A). Cellular localization of L1 (ORF1p) by confocal analysis, suggested that miR-128-induced reductions of nuclear localization of ORF1p was partly, but significantly rescued by overexpressing TNPO1 (WT and miR-128 resistant), as compared to controls (Figure 5B, white arrows indicate examples of ORF1p nuclear staining in the single channel images. Quantification right panels, and Supplemental Figure S6B for WT L1 confocal images). Finally, we analyzed the amount of ORF2 mRNA, as an indirect measure for L1 RNA, in the same experimental conditions, and found that TNPO1 overexpression rescued the miR-128-induced decrease in ORF2 amount, relative to control cells (Supplemental Figure S6C). These results show that overexpression of both WT or miR-128 resistant TNPO1 can partly rescue miR-128-induced L1 restriction.

**Figure 5:**
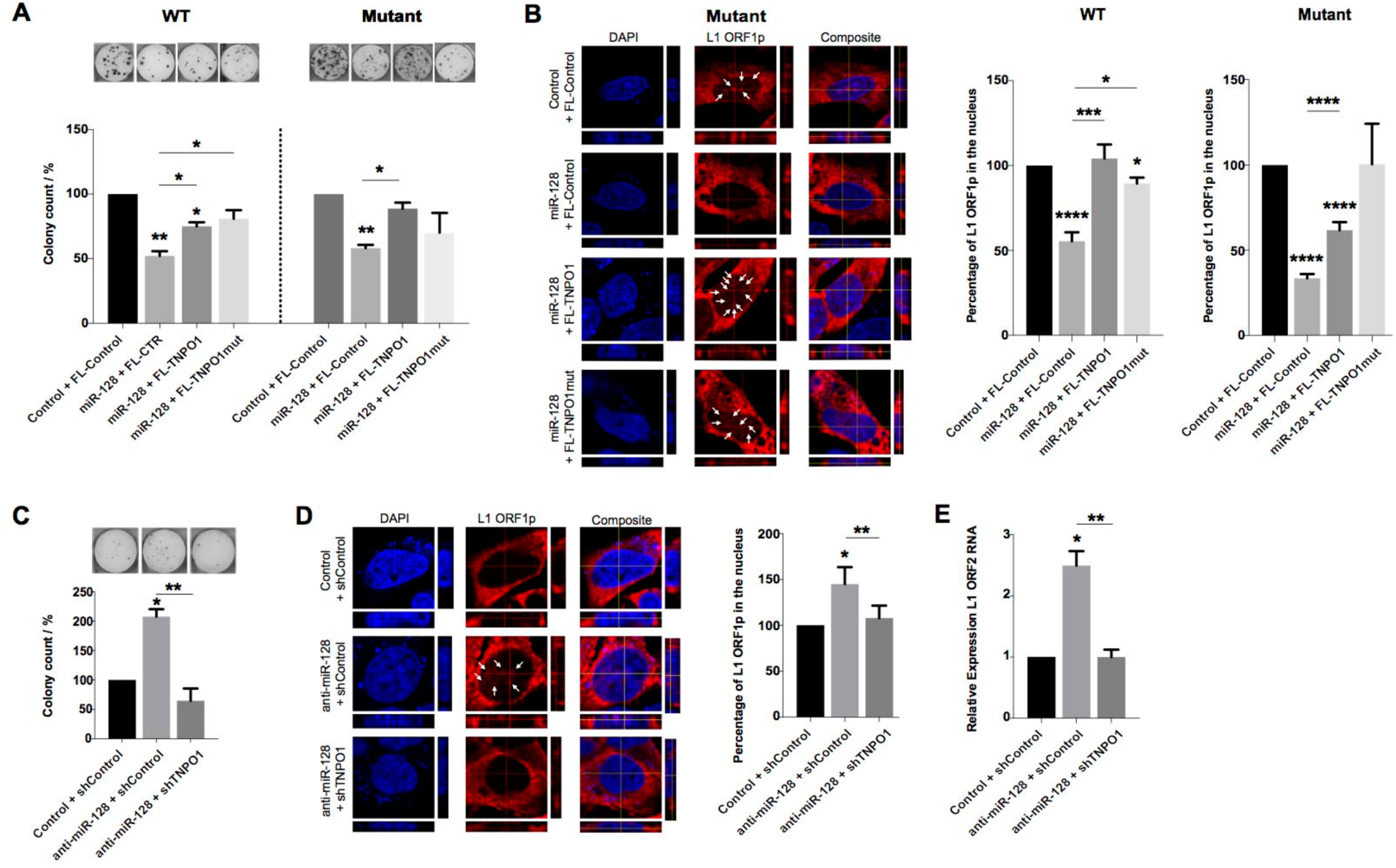
TNPO1 partly rescues miR-128-induced repression of L1 retrotransposition and genomic integration. **(A)** *de novo* retrotranposition was determined by the change in colony count of neomycin-resistant foci of HeLa cells stably transduced with control miR (control) or miR-128, transfected with FL-control, FL-TNPO1 or FL-TNPO1mut (miR-128 resistant), and co-transfected with miR-128 mutant L1 expression plasmid. Results are shown as a mean ± SEM (n=3, independent biological replicates, *, p<0.05, **, p<0.01 and See Supplemental Figure S6A). **(B)** Localization of L1 ORF1p-HA was determined in HeLa cells stably transduced with control miR (control) or miR-128, then transfected with FL-control, FL-TNPO1 or FL-TNPO1mut, and co-transfected with miR-128 mutant L1 expression plasmid. Representative orthogonal views of z-stack images are shown. Quantification of L1 ORF1p-HA localization to the nucleus is shown as the mean percentage of L1 ORF1p in the nucleus (represented as a percentage of L1 ORF1p in nucleus/all L1 ORF1p in the image. White arrows indicate examples of ORF1p nuclear staining in the single channel images) ± SEM (n=50, technical replicates of 3 independent biological replicates, ***, p<0.001, ****, p<0.0001 and See Supplemental Figure S6B). **(C)** New retrotransposition events was determined by change in colony count of neomycin-resistant foci in HeLa cells stably transduced with control miR (control) or anti-miR-128, transfected with shControl or shTNPO1, and co-transfected with wild-type (WT) L1 expression plasmid. Results are shown as a mean ± SEM (n=3, independent biological replicates, *, p<0.05, **, p<0.01). **(D)** Localization of L1 ORF1p-HA was determined in HeLa cells stably transduced with control miR (control) or anti-miR-128, then transfected with shControl or shTNPO1, and co-transfected with wild-type (WT) L1 expression plasmid. Representative orthogonal views of z-stack images are shown. Quantification of L1 ORF1p-HA localization to the nucleus is shown as the mean percentage of L1 ORF1p in the nucleus (White arrows indicate examples of ORF1p nuclear staining in the single channel images) ± SEM (n=50, technical replicates of 3 independent biological replicates, *, p<0.05, **, p<0.01). **(E)** Relative expression of ORF2 amount normalized to B2M in HeLa cells stably transduced with control miR (control) or anti-miR-128, transfected with shControl or shTNPO1, and co-transfected with wild-type L1 plasmid. Results are shown as a mean ± SEM (n=3, independent biological replicates, *, p<0.05, **, p<0.01).

Finally, we performed experiments in which we depleted cell of TNPO1 (using TNPO1 shRNA) in anti-miR-128 stable HeLa cells (in which endogenous levels of miR-128 are neutralized). TNPO1 depletion in anti-miR-128 HeLa cells resulted in a partial and significant rescue of the inhibitory effect of miR-128 on L1 retrotransposition and integration, relative to control, as determined by colony formation assays (Figure 5C), L1 ORF1p nuclear localization by confocal analysis (Figure 5D, white arrows indicate examples of ORF1p nuclear staining in the single channel images, quantification right panel) and amount of ORF2 mRNA (Figure 5E). These combined results strongly support the idea that TNPO1 is a functional target for miR-128 and play an important role in L1 retrotransposition, possibly by affecting nuclear import of L1.

## DISCUSSION

Our present data now provides additional mechanistic context for our earlier report that miRs have adopted part of piRNAs role in somatic cells in order to function as genomic gatekeepers by directly repressing L1 retrotransposon mobilization (37). In addition we show for the first time that a cellular factor (TNPO1) is involved in L1 mobilization by facilitating nuclear import of some L1-RNP complexes and thus gaining access to host DNA. Our results are in alignment with previous reports describing that TNPO1 function in nuclear transport of cargoes including viral proteins (45, 46), and suggests that mobile DNA elements such as L1 elements are part of TNPO1 cargoes. Furthermore, recent data by Marcia et al. demonstrates that L1 efficiently can transposes in non-diving cells (15). We propose that TNPO1 may be involved in active nuclear import of L1-RNP complexes in all cells, but may be crucial for L1 mobilization in non-dividing cells such as neurons. It is possible that TNPO1 functions in a similar fashion during L1-RNP nuclear import as TNPO3 has been demonstrated to assist in nuclear import of the pre-integration complex of HIV-1 (60-66). In summary, we propose a model for miR-128-induced L1 repression in which miR-128 acts by directly targeting L1 RNA (37), as well as indirectly reducing L1 mobilization by repressing a cellular factor involved in nuclear import of some L1-RNP complex (TNPO1) (see Figure 6). We speculate that a dual mechanism helps secure L1 restriction and thus L1-induced retrotransposition and genomic integration in somatic cells.

**Figure 6:**
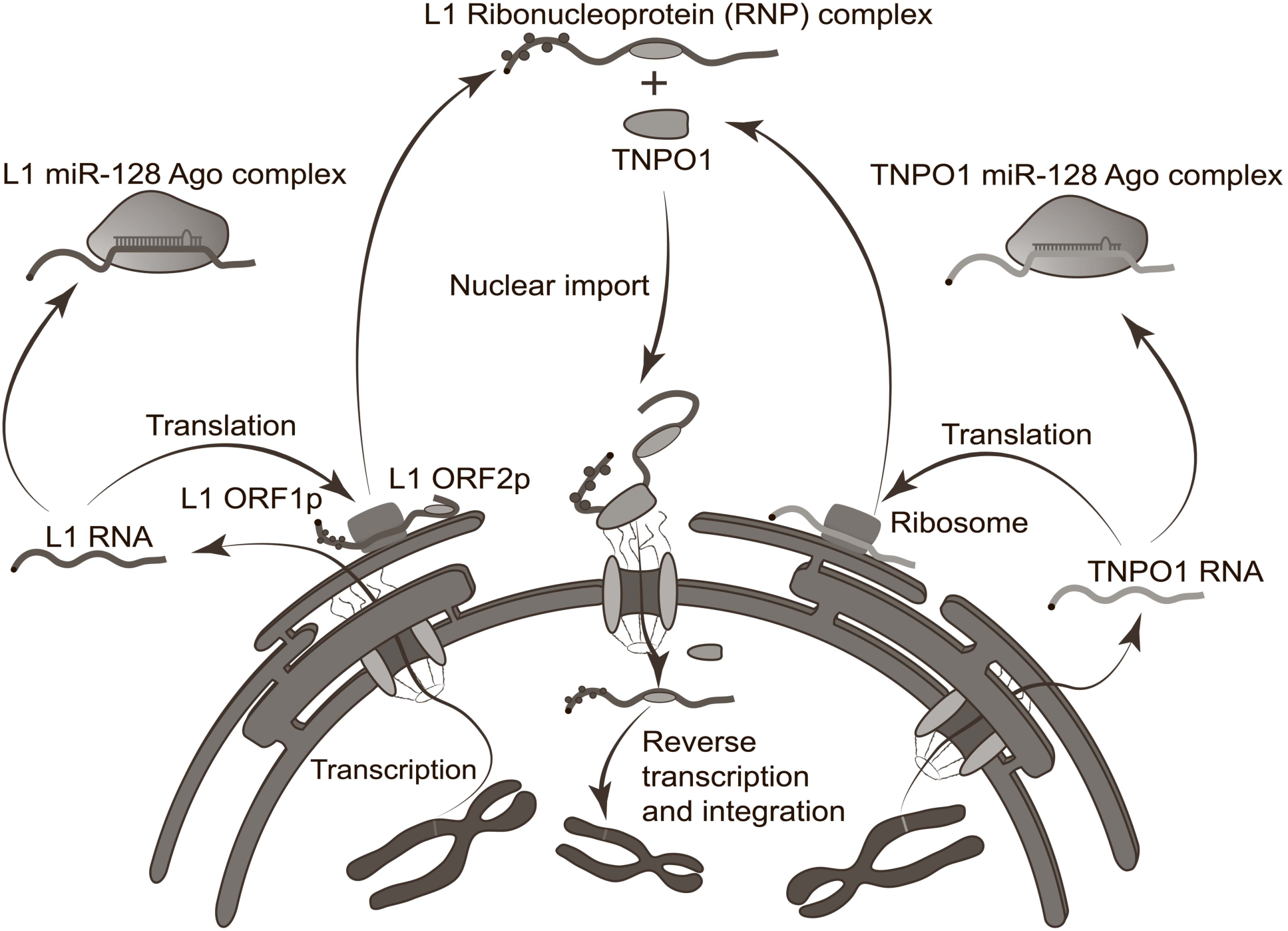
miR-128 regulates L1 retrotransposition by a dual mechanism. Cartoon representation of miR-128-induced repression of L1 retrotransposition and genomic integration. miR-128 inhibits L1 activity by directly targeting L1 RNA, as well as indirectly by repressing the levels of the cellular co-factor TNPO1, which L1 is dependent on for nuclear import and replication.

Interestingly, both TNPO1 and L1 ORF1p have previously, independently been found to interact with the heterogeneous nuclear ribonucleoprotein A1 (hnRNPA1), which contains nuclear localization signals (NLSs) required for shuttling between the cytoplasm and the nucleus (45, 55-58, 68, 71, 72). We have now obtained results, which supports the idea that TNPO1 and ORF1p are also binding partners, either directly or through an RNA bridge (L1 RNA), suggesting a possible scenario in which nuclear import of the L1-RNP complex is assisted through ORF1p, TNPO1 and hnRNPA1 interactions. Another possible scenario is that ORF1p and/or ORF2p could be direct cargoes of TPNO1. While a PY-NLS relies on structure, there is a weak consensus based on characterized motifs (R/H/KX2–5PY). Interestingly, both proteins contain ‘PY’ motifs within the protein sequence, which fits the consensus (perfectly for ORF1p and partly for ORF2p). Future studies will determine whether these motifs are critical for L1 retrotransposition and binding to TPNO1.

The family of TNPO proteins (TNPO1, -2 and TNPO3), all function in nuclear import (44, 55-58, 73, 74). Interestingly, miR-128 harbors predicted binding sites in all three TNPO mRNAs and our preliminary results show that miR-128 down-regulates the expressing level of TNPO1, TNPO2 and TNPO3 mRNAs. This finding has important implications, as TNPO3 is a demonstrated cellular co-factor, which HIV-1 is dependent on for nuclear import of HIV-1 and viral replication (61, 62, 64-66). We anticipate that miR-128-induced TNPO3 repression could have significant effects on the viral life cycle of HIV-1.

Furthermore, miR-128 has previously been demonstrated to function as a tumor suppressor, by inhibition of stemness and epithelial to mesenchymal transition (EMT) through the regulation of target mRNAs including BMI-1, Nanog, HIF-1, VEGF, TGFBR1, EGFR (75-80). We predict, that restriction of L1 insertions is another mechanism by which miR-128 plays a role in inhibiting tumor initiation and tumor cell progression.

Finally, it is interesting to note that the brain expresses ∼70% of all mature miRs, that miR-128 is highly enriched in the brain as compared to other human tissue (81, 82) and that L1 retrotransposition surprisingly have been found derepressed in neuronal progenitor, leading to somatic brain mosaicism and enhanced plasticity (83, 84). These finding suggest a potential important role for miR-128 in the regulation of genomic instability and plasticity in the human brain.

In conclusions, our results show that increased miR-128 expression reduces nuclear import of L1 (ORF1p) and significantly inhibit L1 mobilization; meanwhile, upregulation of TNPO1, a direct and functional target of miR-128, can markedly enhance levels of nuclear L1 (ORF1p) and *de novo* L1 retrotransposition. This newly identified miR-128/TNPO1 module provides a new avenue to an understanding of the L1 life cycle, especially, how some L1-RNP complexes may access host DNA, independently of cell division. Finally, the fact that TNPO1 can partially rescue the miR-128’s inhibitory effect, suggests that miR-128 may repress additional cellular factors, which L1 is dependent on for optimal genomic mobilization.

### Experimental Procedures

#### Cell culture

All cells were cultured at 37°C and 5% CO2. HeLa cells (CCL-2, ATCC) were cultured in EMEM (SH3024401, Hyclone) supplemented with 10% HI-FBS (FB-02, Omega Scientific), 5% Glutamax (35050-061, ThermoFisher), 3% HEPES (15630-080, ThermoFisher), and 1% Normocin (ant-nr-1, Invivogen). Tera-1 cells (HTB-105, ATCC) were cultured in McCoy’s 5A (16600-082, ThermoFisher) supplemented with 20% Cosmic Serum (SH3008702, Fisher Sci), and 1% Normocin (ant-nr-1, Invivogen). 293T cells (CRL-3216, ATCC) were cultured in DMEM supplemented with 10% HI-FBS (FB-02, Omega Scientific), 5% Glutamax (35050-061, Lifetech) and 1% Normocin (ant-nr-1, Invivogen). IMR90-1 cells were cultured in Nutristem complete media (Stemgent). All cell lines were routinely tested for mycoplasma contamination.

#### Transfection and transduction of miRs

OptiMem (31985070, ThermoFisher) and Lipofectamine RNAiMax (13778, ThermoFisher) were used according to manufacturer instructions to complex and transfect 20μM miR-128 mimic or anti-miR-128 (C-301072-01 and IH-301072-02 respectively, Dharmacon) into cells. pJM101/L1 expressing plasmid was co-transfected with miR-128 mimic or anti-miR-128 into cells using OptiMem and Lipofectamine RNAiMax transfection reagent. Optimem and Lipofectamine LTX with plus reagent (15338030, ThermoFisher) were used to complex and transfect 1μg FL-Control or FL-TNPO1 plasmid along with 0.5μg PhiC31 integrase plasmid according to manufacturer instructions. VSVG-pseudotyped lentiviral vectors were made by transfecting (0.67μg of pMD2-G (12259, Addgene), 1.297μg of pCMV-DR8.74 (8455, Addgene), and 2μg of mZIP-miR-128, mZIP-anti-miR-128, pLKO-shControl or pLKO-shTNPO1 (transfer plasmid)) into 293T cells using Lipofectamine LTX with plus reagent (15338030, ThermoFisher). Virus-containing supernatant was collected 48hr and 96hr post-transfection. Viral SUPs were concentrated using PEG-it virus precipitation solution (LV810A-1) according to manufacturer’s instructions. Cells were transduced with high titer virus using polybrene (sc-134220, Santa Cruz Biotech) and spinfection (800g at 32°C for 30 minutes). Transduced cells were then incubated for 48 hours at 37°C and 5% CO2. Cells were selected for 7 days using 3μg/mL Puromycin. Stable lines were maintained in 3μg/mL Puromycin.

#### RNA extraction and quantification of mRNAs

RNA was extracted using Trizol (15596-018, ThermoFisher) and Direct-zol RNA isolation kit (R2070, Zymo Research). cDNA was made with High-Capacity cDNA Reverse Transcription Kit (4368813, ThermoFisher). mRNA levels were analyzed by qRT-PCR using SYBR Green (ThermoFisher) or Forget-me-not qPCR mastermix (Biotium) relative to beta-2-microglobulin (B2m) housekeeping gene and processed using the ΔΔC_t_ method.

#### Immunofluorescence

Cells were fixed with 4% Paraformaldehyde (SigmaAldrich), blocked with 10% Goat serum (ThermoFisher) + 0.1% Triton X-100 (ThermoFisher). Anti-HA antibody was used 1:500 (C29F4, Cell Signaling Technology) and incubated for 24 hours at 4°C. Secondary Goat anti-Rabbit antibody conjugated to a488 (non-GFP expressing cells) or a568 (for GFP expressing cells) was used at 1:500. Slides were mounted with Vectashield containing DAPI counterstain (H-1200, Vector Laboratories). Images were acquired with a Zeiss LSM 700 Confocal microscope in the Optical Biology Core at UC Irvine. Co-localization of ORF1p-HA with the nucleus was calculated as: Percent L1 ORF1p in nucleus = (L1 ORF1p signal co-localized with nucleus/ Total L1 ORF1p signal) X 100. CellProfiler software (47) was utilized to automatically segment nuclei, determine the area of positive staining for L1 ORF1p, the area of positive staining for nuclei (DAPI) and the co-localized area of L1 ORF1p and nuclear staining. Amount of ORF1p in control nuclei was set to 100% and the levels in the experimental nuclei are shown as a percentage of the controls.

#### qPCR screen for additional cellular targets of miR-128

As part of a effort to incorporate authentic research experiences into undergraduate labs at UC Irvine, a basic screen to identify bioinformatically determined cellular targets of miR-128 was performed. Briefly, HeLa cells were transfected with 60 pmol of miR control or miR-128 mimics (GE Dharmacon) using Dharmafect1 (ThermoFisher). After 24 hours, cells were transfected a second time and incubated for another 24 hours after which cells were pelleted and snap frozen in LN2. Control or miR-128 transfected pellets were provided to undergraduate students who isolated RNA and made cDNA using (GeneJET RNA purification kit, ThermoFisher). qPCR was performed using student-designed primers to detect bioinformatically-defined targets of miR-128 (targetscan, pictar). Graduate students further tested differentially expressed targets in independent biological replicates.

#### Site directed mutagenesis

Reverse transcriptase incompetent PJM101/L1 plasmid was made using Q5 Site-directed mutagenesis Kit (E0554S, New England Biolabs) and mutation strategy described in Morrish et al (48) where D702A mutation in L1 ORF2 resulted in an incompetent reverse transcriptase.

#### Cloning

The ORF1-HA gene was generated by polymerase chain reaction (PCR) on DNA from the plamid pJM101/L1 (ORF1). To generate the ORF1-HA insert, we used the sense ORF1-HA primer (5´-GCCTAAGATC TAGGTACCAC CATGGGGAAA AAACAGAACA GAAAAAC-3´) and antisense ORF1-HA primer (5´-GTATCTTATC ATGTCTGGCC AGCTAGC*TTA GGCGTAGTCG GGCACGTCGT AGGGGTAGCC* CATTTTGGCA TGATTTTGCA GCG-3´; HA-tag is shown in italic letters). All amplicons were generated using the Phusion High-Fidelity PCR Kit (NEB) according to the manufacturer´s protocol. The amplicons were cloned into the expression vector pExpress-mUKG-MH1 by replacing the mUKG insert by the amplicon. For the generation of the plasmid backbone the vector was cut by *Nco*I and *Nhe*I and the insert was cloned into the backbone by using the cold fusion kit (SBI) according to the manufacturer´s protocol. The resulting plasmid pExpress-ORF1-HA-MH1, pExpress-ORF1-HA-MH1 was amplified in *E. coli* and validated by sequencing.

TNPO1 shRNA was designed using the RNAi Consortium (https://www.broadinstitute.org/rnai/public/) using clone TRCN0000382164 and cloned into pLKO.1 puro backbone (Addgene, #8453). pLKO shGFP control plasmid was pre-assembled (Addgene, #30323).

For the TNPO1 full-length clone we modified the plasmid pFC-PGK-MCS-pA-EF1-GFP-T2A-Puro (SBI, backbone) by replacing the PGK with a CMV promoter. The CMV promoter provides a strong and robust expression on most cell types. The CMV promoter was amplified by PCR from the phiC31 integrase expression plasmid (SBI). To generate the CMV promoter insert, we used the sense CMV primer (5´-CTAGAACTAG TTATTAATAG TAATCAATTA CGGGGTC-3´) and antisense CMV primer (5´-GATATCGGAT CCACCGGTAC CAAGCTTAAG TTTAAAC-3´). The insert and the backbone of the plasmid were cut by *Xba*I and *BamH*I and purified by an agarose gel. Insert and backbone were ligated together using the quick ligation kit (NEB) and transformed. The resulting plasmid pFC-CMV-MCS-pA-EF-1-GFP-T2A-Puro-MH1 was verified by sequencing.

For the cloning of the full-length TNPO1 mRNA expression clone (FL-TNPO1), we isolated total RNA from A549 and HeLa cells. 20 ng of the total RNA was reverse transcribed using a poly dT primer. For the amplification of the TNPO1 gene, we used the sense TNPO1 sense (5´-TTTAAACTTA AGCTTGGTAC CGGTGGATCC GCCACCATGG AGTATGAGTG GAAACCTGAC-3´) and antisense TNPO1 antisense (5´-GATTAAACAC CATAAAAAGC TGCA -3´). The 3´-UTR of the gene that exhibits the binding site for miR-128 was split into four fragments. For the four parts, the following primers were used: part I 3´-UTR primer sense 1 (5´-GGAAGGGTAA ACCAGTAGGG AATA -3´) and 3´-UTR antisense 1 (5´-GGGTTAACTT AACAAGGATT TATTCAC-3´); part II 3´-UTR primer sense 2 (5´-CTGTGAATAA ATCCTTGTTA AGTTAAC-3´) and 3´-UTR antisense 2 (5´-GTAAACACTG ACCTCCTGAG GTTCCTA-3´); part III 3´-UTR primer sense 3 (5´-GTAGGAACCT CAGGAGGTCA GTGTTTA-3´) and 3´-UTR antisense 3 (5´-GGGATACAAA CCACAATGAACAAT-3´), part IV 3´-UTR primer sense 4 (5´-CAATTGTTCA TTGTGGTTTG TATC-3´) and (5´-GGCAACTAGA AGGCACAGTC GATCGATTAT AGTTAAACAA CTTTATTAACATAGTCAAGC-3´). All amplicons were generated using the Phusion High-Fidelity PCR Kit (NEB) according to the manufacturer´s protocol. The fragments were stepwise assembled by using the cold fusion kit (SBI) and cloned into the pFC-CMV-MCS-pA-EF-1-GFP-T2A-Puro-MH1 *Bam*HI/*Cla*I linearized backbone by cold fusion. The resulting plasmid pFC-CMV-TNPO1-pA-EF-1-GFP-T2A-Puro-MH1 was verified by sequencing. FL-Control is an empty vector.

#### Colony formation assay

Stable HeLa lines expressing miR control, miR-128, anti-miR-128, shControl, shTNPO1, FL-Control, FL-TNPO1 were plated at 5x10^5^ cells per well of a 6-well plate and incubated for 24 hours. Cells were then transfected with 0.5μg of pJM101/L1RP or pJM101/L1RP RT-(containing neomycin resistance retrotransposition indicator cassette) per well using X-treme gene HP DNA transfection reagent (06366236001, Roche) according to manufacturer instructions. Cells were incubated for 24 hours followed by a media change without antibiotics. 48 hours after transfection, selection by daily media changes containing 500μg/mL G418 (ant-gn-1, Invivogen) were initiated. Daily media changes were continued until all cells died in the negative control (untransfected HeLa cells). Neomycin-resistant colonies were fixed with cold 1:1 methanol:acetone and then visualized using May-Grunwald (ES-3410, ThermoFisher) and Jenner-Giemsa staining kits (ES-8150, ThermoFisher) according to manufacturer protocol.

#### Luciferase assays

Wild-type (WT) TNPO1 binding site 1, WT TNPO1 binding site 2, WT TNPO1 binding site 3, mutated TNPO1 binding site 1 or positive control (See supplemental for sequences) sequences were cloned into dual-luciferase reporter plasmid (pEZX-MT05, Genecopoeia). 3x10^5^ HeLa cells were transfected at the same time as seeding with 0.8μg of reporter plasmid (WT, mutated, Pos), and 20nM miR-128 mimic (Dharmacon) or Control mimic (Dharmacon) using Attractene transfection reagent (301005, Qiagen) according to manufacturer instructions. Cells were incubated for 48 hours at 37°C at 5% CO2 without media change. Relative Gaussia Luciferase and secreted alkaline phosphatase (SEAP) was determined using Secrete-Pair Dual Luminescence Assay Kit (SPDA-D010, Genecopoeia) in technical triplicates from collected supernatant. Luminescence was detected by Tecan Infinite F200 Pro microplate reader.

#### Fractionation

Nuclear and cytoplasmic fractions were isolated using Protein and RNA Isolation System or PARIS kit (AM1921, ThermoFisher) according to manufacturer instructions 48 hours after transfection. Both RNA and protein were isolated from the same biological sample (shControl, shTNPO1, FL-Control, FL-TNPO1) and used for qRT-PCR and corresponding Western blot analysis respectively.

#### Immunoprecipitation

Transfected cells were lysed in RIPA buffer + Protease inhibitors on ice for 15minutes. Lysate was cleared by centrifuging at 13,000 rpm (max speed tabletop centrifuge) for 15minutes at 4°C. Supernatant was collected and a portion was reserved as “INPUT” (IN) for Western analysis. Remaining IP was mixed with protein G beads (NEB) and 5μg of anti-HA antibody and incubated on a rotator for 24 hours at 4°C. Beads were separated using a magnetic rack and washed four times with PBS. Beads were then boiled for 5 minutes at 95°C in 4X Protein Loading Dye (+SDS). Beads were separated and supernatant containing the immunoprecipitated proteins was used for Western analysis.

#### Western blot

Mouse anti-human L1 ORF1p (MABC1152 clone 4H1) from EMD Millipore was used at 1:1000, alternatively Rabbit anti-human L1 ORF1p antibody custom generated by Genscript against and validated by ELISA was used at 1:1000. Western blot analysis of Genscript antibody was initially cross-checked by a custom generated anti-human L1 ORF1p antibody (kindly provided by G. Schumann). Rabbit anti-HA antibody to detect HA-tagged L1 ORF1p (C29F4, Cell Signaling Technology) was used at 1:5000, Mouse anti-TNPO1 antibody was used at 1:2000 (ab10303, Abcam), Rabbit anti-hnRNPA1 antibody was used at 1:2000 (K350, Cell Signaling Technology), anti-alpha Tubulin antibody (ab4074, Abcam) was diluted 1:5000, and anti-GAPDH antibody (14C10, Cell Signal Technology) diluted 1:3000 were used as loading controls, validation can be found on the manufacturer websites. Secondary HRP-conjugated anti-rat (ab102172, Abcam), HRP-conjugated anti-rabbit (GE), and HRP-conjugated anti-mouse (GE) was used at 1:5000. ECL substrate (32106, ThermoFisher) was added and visualized on a BioRad ChemiDoc imager. Since many proteins were similar in size, blots were not cut but developed sequentially. After developing, each blot was washed three times in 1X TBST (TBS, #BP24711, FisherSci + 0.05% Tween 20, #BP337-500, FisherSci) prior to incubation with the next primary antibody.

#### Argonaute RNA Immunopurifications (Ago RIP)

Immunopurification of Argonaute from HeLa cell extracts was performed using the 4F9 antibody [4F9, Santa Cruz Biotechnology] as described previously (49, 50). Briefly, 10mm plates of 80% confluent cultured cells were washed with buffer A [20 mM Tris-HCl pH 8.0, 140 mM KCl and 5 mM EDTA] and lysed in 200ul of buffer 2XB [40 mM Tris-HCl pH 8.0, 280 mM KCl, 10 mM EDTA, 1% NP-40, 0.2% Deoxycholate, 2X Halt protease inhibitor cocktail (Pierce), 200 U/ml RNaseout (ThermoFisher) and 1 mM DTT. Protein concentration was adjusted across samples with buffer B [20 mM Tris-HCl pH 8.0, 140 mM KCl, 5 mM EDTA pH 8.0, 0.5% NP-40, 0.1% deoxycholate, 100 U/ml Rnaseout (ThermoFisher), 1 mM DTT and 1X Halt protease inhibitor cocktail (Pierce)]. Lysates were centrifuged at 16,000g for 15 mins at 4^o^C and supernatants were incubated with 10-20 ug of 4F9 antibody conjugated to epoxy magnetic beads (M-270 Dynalbeads, ThermoFisher) for 2 hours at 4^o^C with gentle rotation (Nutator). The beads, following magnetic separation, were washed three times five mins with 2 ml of buffer C [20 mM Tris-HCl pH 8.0, 140 mM KCl, 5 mM EDTA pH 8.0, 40 U/ml Rnaseout (ThermoFisher), 1 mM DTT and 1X Halt protease inhibitor cocktail (Pierce)]. Following immunopurification, RNA was extracted using miRNeasy kits (QIAGEN), following the manufacturer’s recommendations and qPCR was performed using custom probes/primers for the TNPO1 mRNA transcript and Forget-me-not qPCR mastermix (Biotium). Results were normalized to their inputs and shown as “corrected” values as a proxy for Ago immunopurification efficiency

#### Statistical analysis

Student’s t-tests were used to calculate two-tailed *p* values and data are displayed as mean ± standard error of the mean (SEM) of technical (TR) or independent biological replicates (IBR), (n) as indicated.

## ACKNOWLEDGEMENTS

We thank the undergraduate students involved in the miR-128 screen effort, as part of laboratory class at University of California, Irvine (UCI), and Y. Castuera and D. Jury for lab assistance in quantifying results. We also thank J. Moran (University of Michigan Medical School, Ann Arbor, MI) and M. An (Washington State University, Pullman, WA) for generously sharing pJM101/L1 and pWA355 plasmids. And finally we thank A. Mortazavi (University of California, Irvine) for generously sharing of reagents and support.

## AUTHOR CONTRIBUTION

A. Idica performed the majority of experiments demonstrating that miR-128 targets TNPO1 and that miR-128-induced L1 restriction is partly dependent on TNPO1. A. Idica also helped generate the first version of the figures and gave input on the manuscript. EA. Sevrioukov performed colony formation assays, and cross-validated ORF1p antibodies. D. Zisoulis performed the Ago RNA IPs. M. Hamdorf generated the miR-128 resistant L1 and TNPO1-overexpressing plasmids. I. Daugaard generated the final figures and commented on the manuscript. P. Kadandale directed the evaluation of miR-128 targets in the lab class module and IM. Pedersen performed some experiments, directed all experiments, figure design and wrote the manuscript. All authors reviewed the results and approved the final version of the manuscript.

---

**Figure S1:**
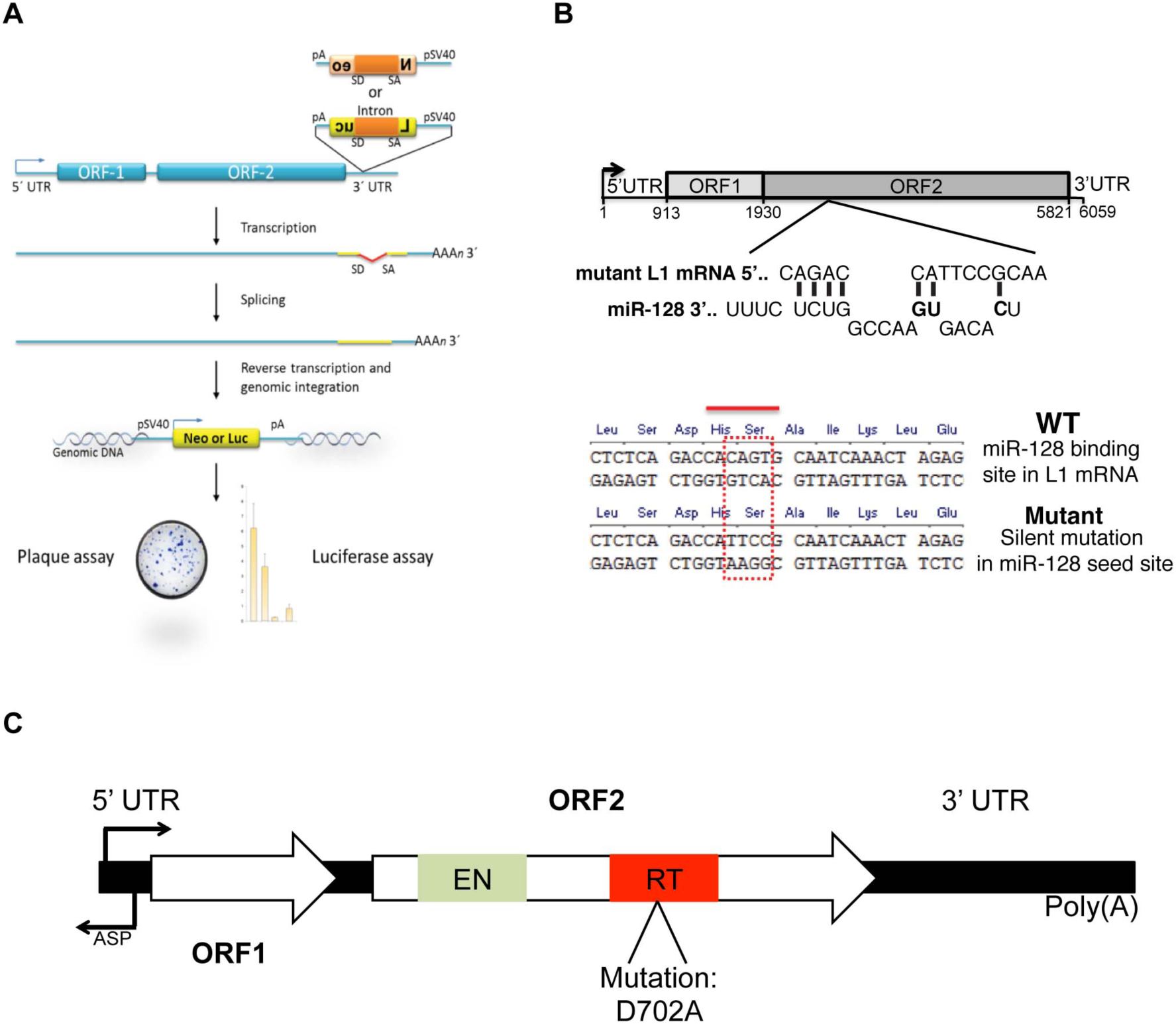
(A) Schematic of the luciferase or neomycin-resistance based LINE-1retrotransposition reporter: a retrotransposition indicator cassette, consisting of a SV40 driven neomycin-resistance or a firefly luciferase gene is inserted in antisense orientation into the 3’UTR of a fulllength L1 element and disrupted by an intron in the sense orientation. The resistance marker or luciferase can only be activated after one round of L1 retrotransposition, when the spliced variant is integrated into the genome, thus allowing the selection or quantification of luciferase of cells with new retrotransposition events in culture. (C) Diagram of miR-128 binding site in ORF2 of L1 mRNA. The mutant plasmid (mutated seed sequence, miR-128 mutant) encodes the full-length L1 mRNA with a silent mutation in the protein coding sequence but introduced a mismatch in the seed sequence-binding site for miR-128. The plasmid encodes a fully functional protein and LINE-1 activity, but does not serve as a perfect miR-128 binding site (shown in red). (B) Schematic following the strategy of Wei et al, a O702A mutation was introduced in the RT domain of L1 ORF2 in the WT plasmid (pJM101) generated a retrotransposon-deficient copy of L1, allowing forthe exclusion of exogenous L1 activity in RT dead experiments.

**Figure S2:**
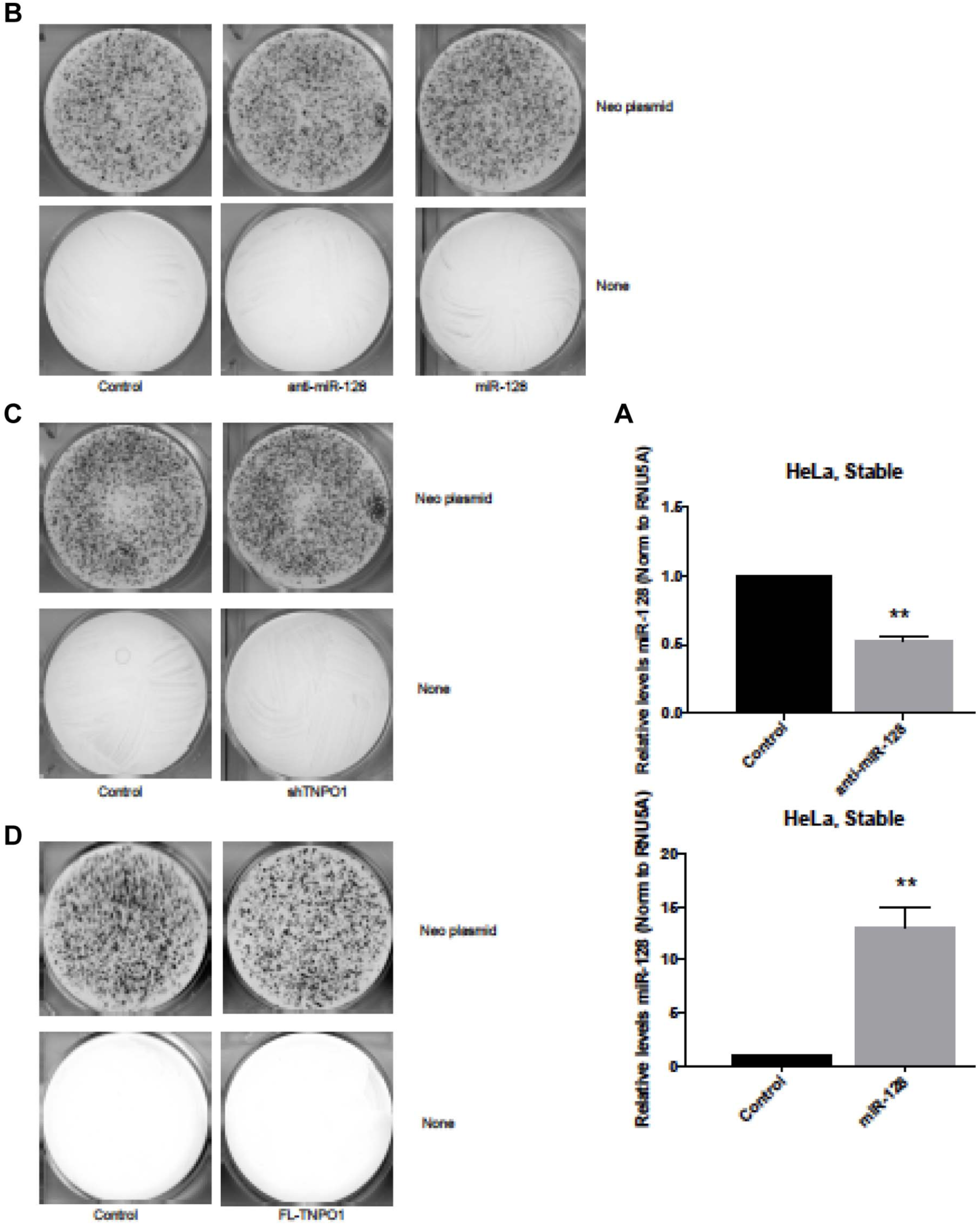
(A) miR-128 expression levels in miR-128-overexpressing and knock down JHela cells, determined by miR-128 specific RT and qPCR analysis. (B) Hela plasmid levels (Neo plasmid, constitutive Neomycin-resistance only) control for Colony Formation Assay verified that miR-128 modulation itself did not affect plasmid transfection levels (neo resistant colonies) as compared to cells non-modulated controls. (C) Knockdown of TNPOl using shRNA did not affect plasmid transfection levels for Colony Formation Assay as shown by neomycinresistant clones. (D) Overexpression of TNPOl did not affect plasmid transfection levels for Colony Formation Assay as shown by neomycin-resistant clones.

**Figure S3:**
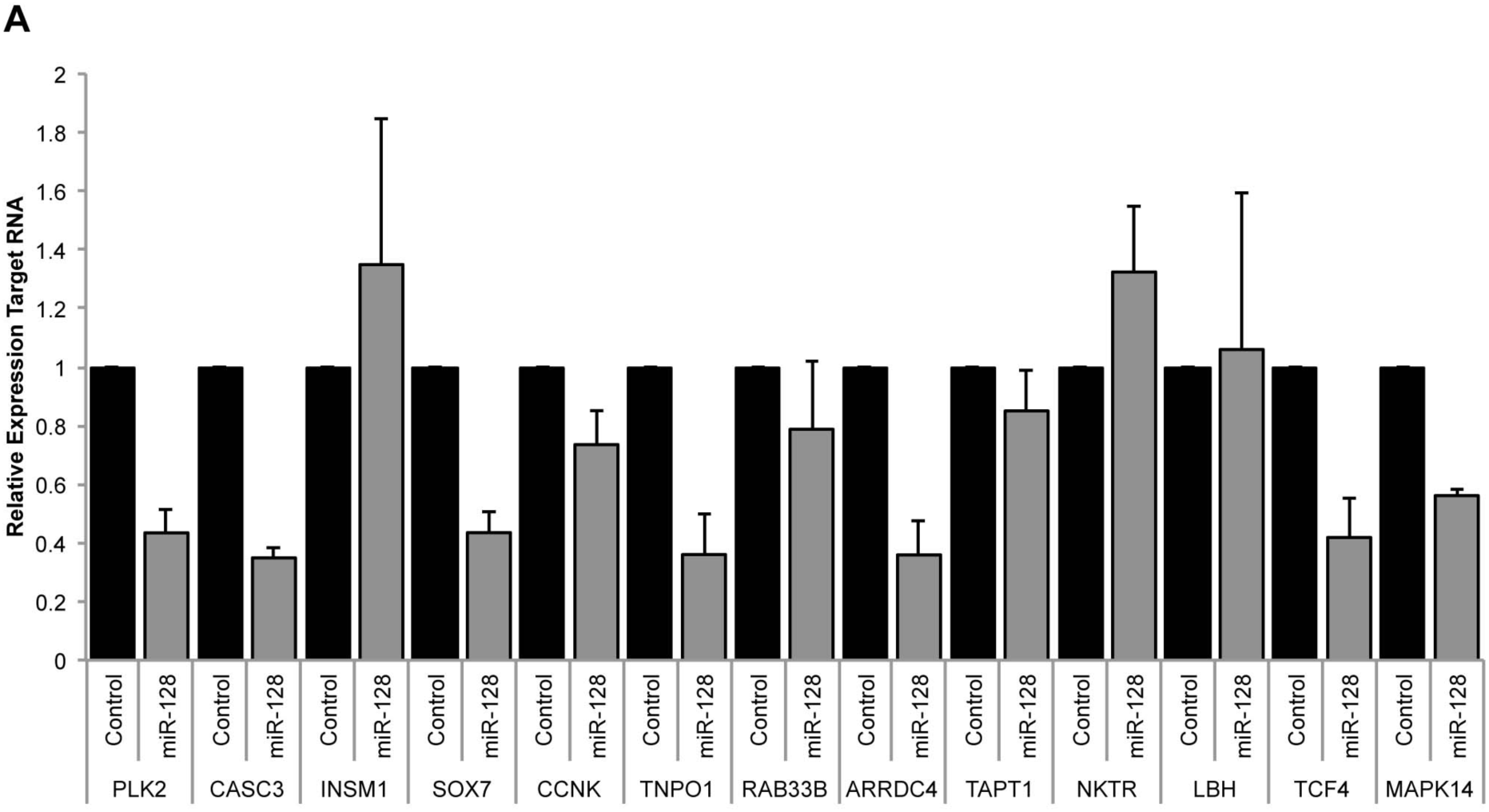
Experimentally-validated bioinformatic targets of miR-128. Relative levels of target RNA normalized to B2m from Hela cells transiently transfected with miR-control mimic or miR-128 mimic were determined for 13 bioinformatically-determined targets.

**Figure S4:**
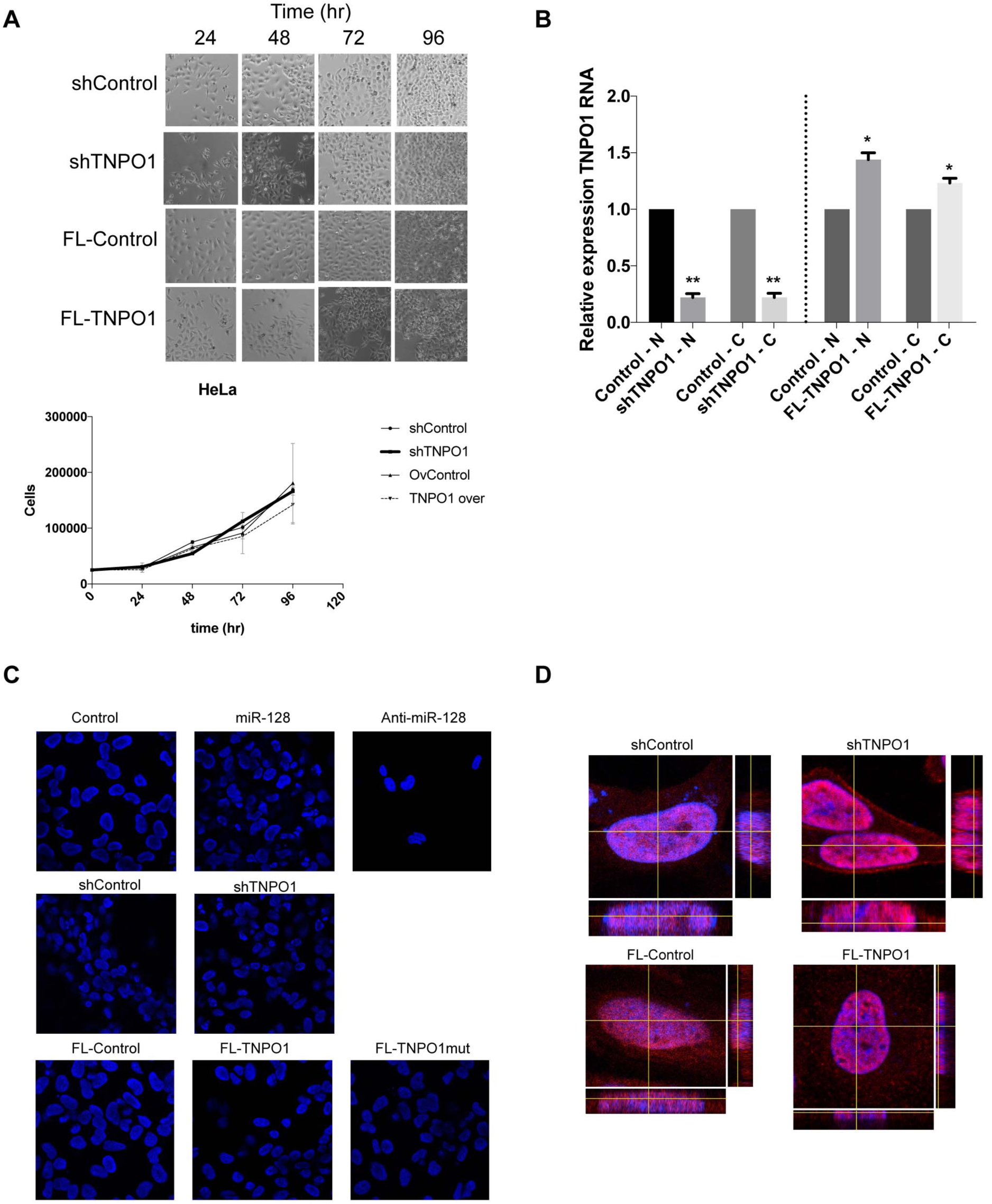
TNP01 controls. (A) TNP01 modualted Hela cells were analyzed for toxicity and effects on cell proliferation. (B) TNP01 mRNA levels was determined by qPCR analysis in nuclear and cyto plasmic fractions of Hela cells. (C) TNP01 and miR-128 modulated cells were stained with Dapi prior to L1 transfections to verify that cells were not contaminated. (D) TBP associated factor 15 (TAB15), a verified TNP01 cargo was used as a positive control, showing thatTNP01 modulation inhibits versus enhances levels of nuclear TAB15.

**Figure S5:**
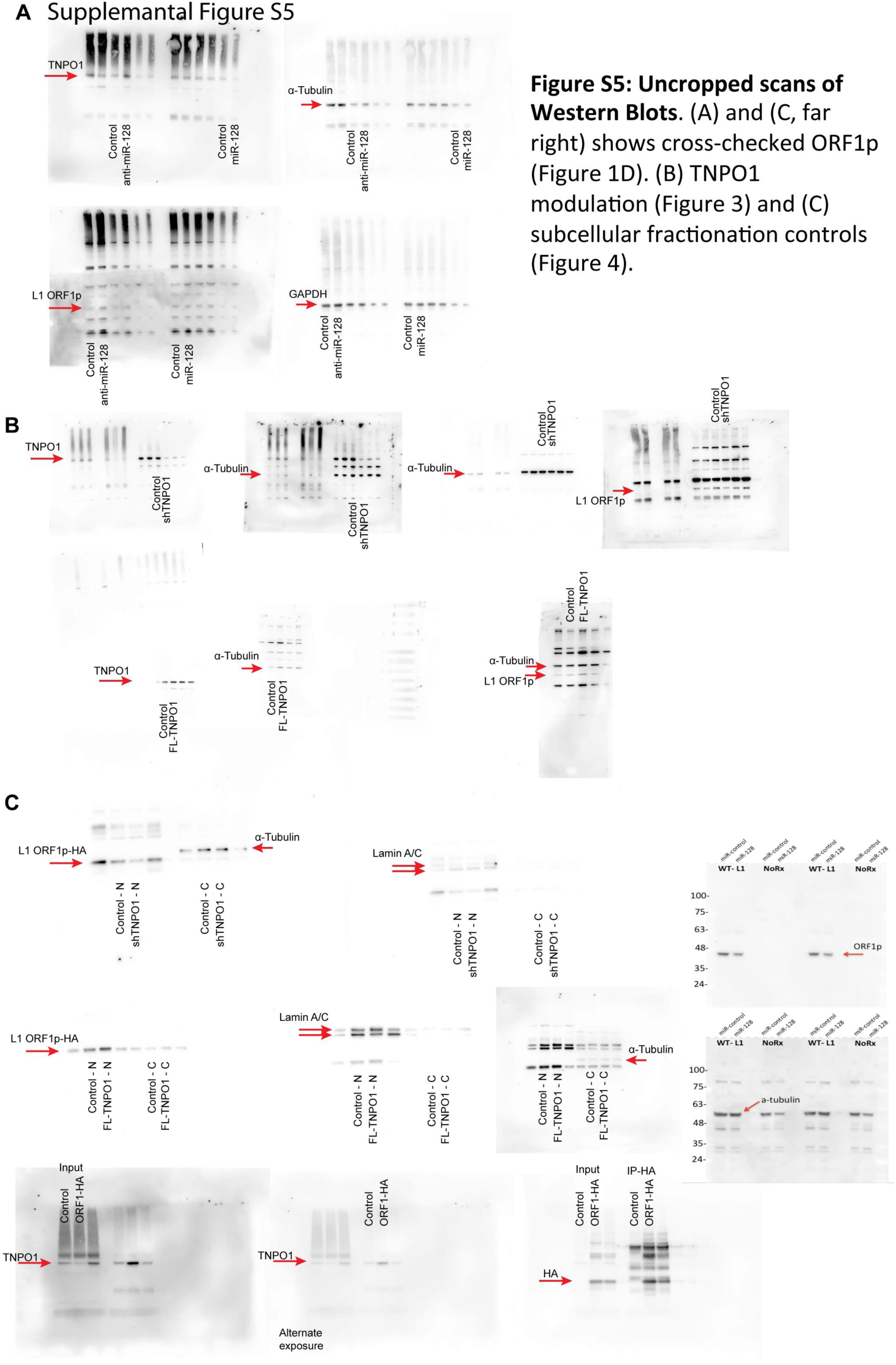
Uncropped scans of Western Blots. (A) and (C, far right) shows cross-checked ORFlp (Figure ID). (B) TNPOl modulation (Figure 3) and (C) subcellular fractionation controls (Figure 4).

**Figure S6:**
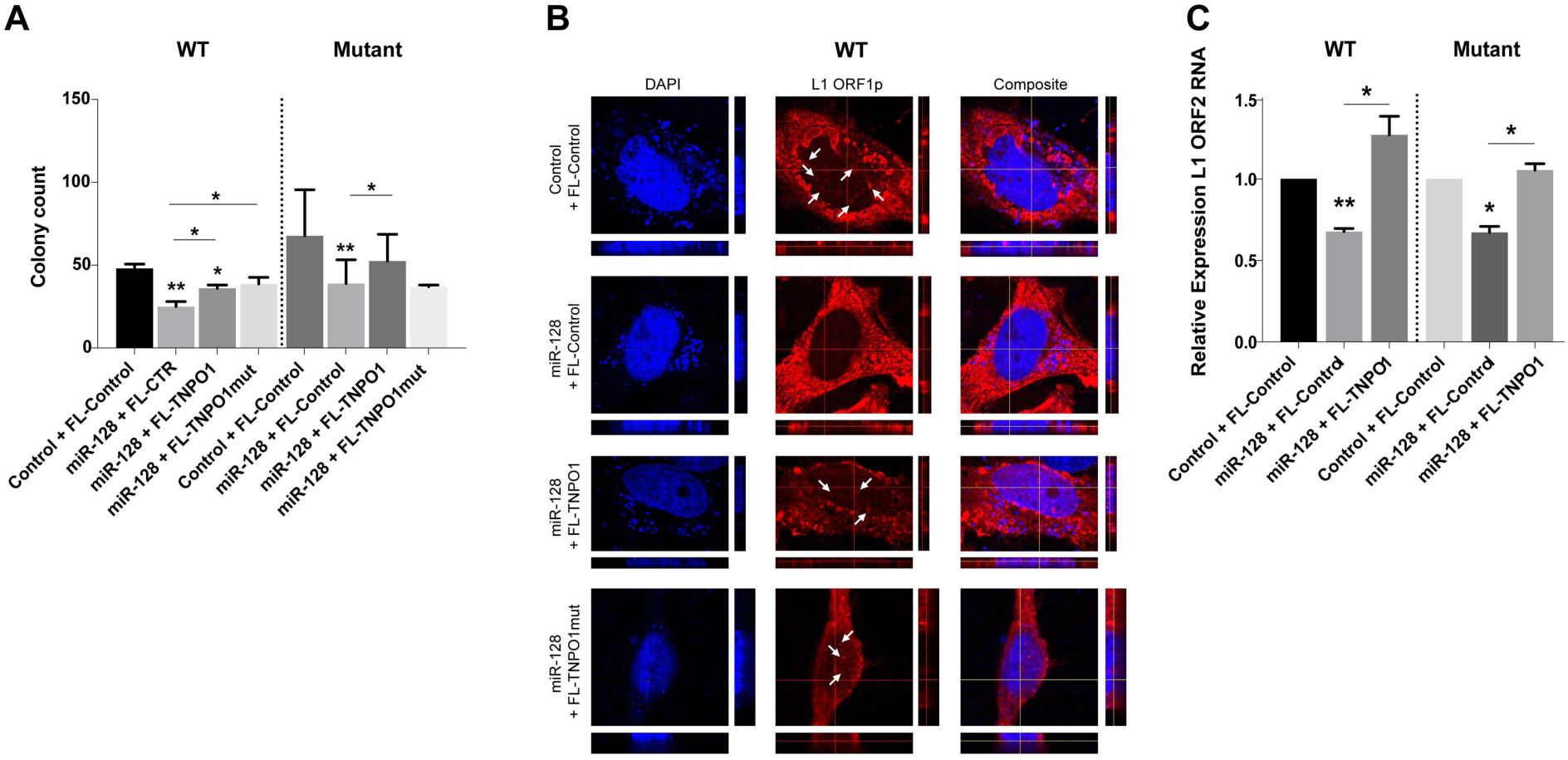
(A) HeLA cells over-expressing plasmid control (FL-CTL), FL-TNPOl or miR-128 resistantTNPOl (FL-TNPOl mut) were transfected with the either a wild-type (WT) or a miR-128 resistant L1 (Mutant) Ll reporter and colony formation assays were performed (shown asco lony counts). (B)Th e same cell lines described in were transfected with WT L1 and OR Fl p-HA and ORFl p cellular localization was analyzed by confocal analysis. White arrows highlights ORFl p expression in the single channel middle pictures. (C) The same cell lines were analyzed for ORF2 mRNA amount by qPCR analysis, normalized to B2M mRNA levels.

## REFERENCES

Lander, E.S., Linton, L.M., Birren, B., Nusbaum, C., Zody, M.C., Baldwin, J., Devon, K., Dewar, K., Doyle, M., FitzHugh, W., et al. (2001) Initial sequencing and analysis of the human genome. Nature, 409, 860–921.

Moran, J.V., Holmes, S.E., Naas, T.P., DeBerardinis, R.J., Boeke, J.D. and Kazazian, H.H., Jr. (1996) High Frequency Retrotransposition in Cultured Mammalian Cells. Cell, 87, 917–927.

Scott, A.F., Schmeckpeper, B.J., Abdelrazik, M., Comey, C.T., O’Hara, B., Rossiter, J.P., Cooley, T., Heath, P., Smith, K.D. and Margolet, L. (1987) Origin of the human L1 elements: proposed progenitor genes deduced from a consensus DNA sequence. Genomics, 1, 113–125.

Speek, M. (2001) Antisense promoter of human L1 retrotransposon drives transcription of adjacent cellular genes. Mol Cell Biol, 21, 1973–1985.

Swergold, G.D.G. (1990) Identification, characterization, and cell specificity of a human LINE-1 promoter. Mol Cell Biol, 10, 6718–6729.

Yang, N. and Kazazian, H.H. (2006) L1 retrotransposition is suppressed by endogenously encoded small interfering RNAs in human cultured cells. Nat Struct Mol Biol, 13, 763–771.

Kolosha, V.O. and Martin, S.L. (1997) In vitro properties of the first ORF protein from mouse LINE-1 support its role in ribonucleoprotein particle formation during retrotransposition. Proc Natl Acad Sci U S A, 94, 10155–10160.

Feng, Q., Moran, J.V., Kazazian, H.H., Jr. and Boeke, J.D. (1996) Human L1 Retrotransposon Encodes a Conserved Endonuclease Required for Retrotransposition. Cell, 87, 905–916.

Mathias, S.L., Scott, A.F., Kazazian, H.H., Boeke, J.D. and Gabriel, A. (1991) Reverse transcriptase encoded by a human transposable element. Science, 254, 1808–1810.

Kleckner, N. (1990) Regulation of transposition in bacteria.-PubMed-NCBI. Annu. Rev. Cell. Biol., 6, 297–327.

Luan, D.D., Korman, M.H., Jakubczak, J.L. and Eickbush, T.H. (1993) Reverse transcription of R2Bm RNA is primed by a nick at the chromosomal target site: a mechanism for non-LTR retrotransposition. Cell, 72, 595–605.

Kubo, S., Seleme, MC., Soifer, HS., Perez, JL., Moran, JV., Kazazian, HH., Jr, Kasahara N. (2006) L1 retrotransposition in nondividing and primary human somatic cells. Proc Natl Acad Sci U S A. 103:8036–41.

Shi, X., Seluanov, A., Gorbunova, V. (2007) Cell divisions are required for L1 retrotransposition. Mol Cell Biol. 27:1264–70.

Xie, Y., Mates, L., Ivics, Z., Izsvák, Z., Martin, S.L, An, W. (2013) Cell division promotes efficient retrotransposition in a stable L1 reporter cell line. Mob DNA. 4:10.

Macia, A., Widmann, TJ., Heras, SR., Ayllon, V., Sanchez, L., Benkaddour-Boumzaouad, M., Muñoz-Lopez, M., Rubio, A., Amador-Cubero, S., Blanco-Jimenez, E., Garcia-Castro, J., Menendez, P., Ng, P., Muotri, AR., Goodier, JL., Garcia-Perez, JL. (2017) Engineered LINE-1 retrotransposition in nondividing human neurons. Genome Res. 27:335–348.

Nigumann, P., Redik, K., Mätlik, K. and Speek, M. (2002) Many Human Genes Are Transcribed from the Antisense Promoter of L1 Retrotransposon. Genomics, 79, 628–634.

Perepelitsa-Belancio, V. and Deininger, P. (2003) RNA truncation by premature polyadenylation attenuates human mobile element activity. Nat Genet, 35, 363–366.

Moran, J.V., DeBerardinis, R.J. and Kazazian, H.H. (1999) Exon Shuffling by L1 Retrotransposition. Science, 283, 1530–1534.

Kazazian, H.H. (2004) Mobile elements: drivers of genome evolution. Science, 303, 1626–1632.

Hancks, D.C. and Kazazian, H.H., Jr. (2012) Active human retrotransposons: variation and disease. Curr. Opin. Genet. Dev., 22, 191–203.

Beck, C.R., Garcia-Perez, J.L., Badge, R.M. and Moran, J.V. (2011) LINE-1 Elements in Structural Variation and Disease. Annu. Rev. Genom. Human Genet., 12, 187–215.

Cordaux, R. and Batzer, M.A. (2009) The impact of retrotransposons on human genome evolution. Nat Rev Genet, 10, 691–703.

Shukla, R., Upton, K.R., Muñoz-Lopez, M., Gerhardt, D.J., Fisher, M.E., Nguyen, T., Brennan, P.M., Baillie, J.K., Collino, A., Ghisletti, S., et al. (2013) Endogenous retrotransposition activates oncogenic pathways in hepatocellular carcinoma. Cell, 153, 101–111.

Aravin, A.A., Sachidanandam, R., Girard, A., Fejes-Toth, K. and Hannon, G.J. (2007) Developmentally Regulated piRNA Clusters Implicate MILI in Transposon Control. Science, 316, 744–747.

Kuramochi-Miyagawa, S., Watanabe, T., Gotoh, K., Totoki, Y., Toyoda, A., Ikawa, M., Asada, N., Kojima, K., Yamaguchi, Y., Ijiri, T.W., et al. (2008) DNA methylation of retrotransposon genes is regulated by Piwi family members MILI and MIWI2 in murine fetal testes. Genes & Development, 22, 908–917.

Tsutsumi, Y. (2000) Hypomethylation of the retrotransposon LINE-1 in malignancy. Jpn. J. Clin. Oncol., 30, 289–290.

Smith, Z.D., Chan, M.M., Mikkelsen, T.S., Gu, H., Gnirke, A., Regev, A. and Meissner, A. (2012) A unique regulatory phase of DNA methylation in the early mammalian embryo. Nature, 484, 339–344.

Chalitchagorn, K., Shuangshoti, S., Hourpai, N., Kongruttanachok, N., Tangkijvanich, P., Thong-ngam, D., Voravud, N., Sriuranpong, V. and Mutirangura, A. (2004) Distinctive pattern of LINE-1 methylation level in normal tissues and the association with carcinogenesis.-PubMed-NCBI. Oncogene, 23, 8841–8846.

Wissing, S.S., Muñoz-Lopez, M.M., Macia, A.A., Yang, Z.Z., Montano, M.M., Collins, W.W., Garcia-Perez, J.L.J., Moran, J.V.J. and Greene, W.C.W. (2012) Reprogramming somatic cells into iPS cells activates LINE-1 retroelement mobility. Hum Mol Genet, 21, 208–218.

Koito, A. and Ikeda, T. (2014) Intrinsic restriction activity by AID/APOBEC family of enzymes against the mobility of retroelements. Mobile Genetic Elements, 1, 197–202.

Bogerd, H.P., Wiegand, H.L., Doehle, B.P. and Cullen, B.R. (2007) The intrinsic antiretroviral factor APOBEC3B contains two enzymatically active cytidine deaminase domains. Virology, 364, 486–493.

Ambros, V. (2004) The functions of animal microRNAs. Nature, 431, 350–355.

Heras, SR., Macias, S., Plass, M., Fernandez, N., Cano, D., Eyras, E., Garcia-Perez, JL., Cáceres, JF. (2013) The Microprocessor controls the activity of mammalian retrotransposons. Nat Struct Mol Biol. 20:1173–81.

Bartel, D. (2004) MicroRNAs: Genomics, Biogenesis, Mechanism, and Function. Cell, 116, 281–297.

Bartel, D.P. (2009) MicroRNAs: Target Recognition and Regulatory Functions. Cell, 136, 215–233.

Fabian, M.R., Sonenberg, N. and Filipowicz, W. (2010) Regulation of mRNA Translation and Stability by microRNAs. Annual Review of Biochemistry, 79, 351–379.

Hamdorf, M., Idica, A., Zisoulis, D.G., Gamelin, L., Martin, C., Sanders, K.J. and Pedersen, I.M. (2015) miR-128 represses L1 retrotransposition by binding directly to L1 RNA. Nat Struct Mol Biol, 22, 824–831.

Vella, M.C., Choi, E.-Y., Lin, S.-Y., Reinert, K. and Slack, F.J. (2004) The C. elegans microRNA let-7 binds to imperfect let-7 complementary sites from the lin-41 3’UTR. Genes & Development, 18, 132–137.

Lin, C.-W., Chang, Y.-L., Chang, Y.-C., Lin, J.-C., Chen, C.-C., Pan, S.-H., Wu, C.-T., Chen, H.-Y., Yang, S.-C., Hong, T.-M., et al. (2013) MicroRNA-135b promotes lung cancer metastasis by regulating multiple targets in the Hippo pathway and LZTS1. Nat Comms, 4, 1877.

Kent, O.A., Fox-Talbot, K. and Halushka, M.K. (2012) RREB1 repressed miR-143/145 modulates KRAS signaling through downregulation of multiple targets. Oncogene, 32, 2576–2585.

Le, M.T.N., Xie, H., Zhou, B., Chia, P.H., Rizk, P., Um, M., Udolph, G., Yang, H., Lim, B. and Lodish, H.F. (2009) MicroRNA-125b promotes neuronal differentiation in human cells by repressing multiple targets.-PubMed-NCBI. Mol Cell Biol, 29, 5290–5305.

Lee, B.J., Cansizoglu, A.E., Süel, K.E., Louis, T.H., Zhang, Z. and Chook, Y.M. (2006) Rules for nuclear localization sequence recognition by karyopherin beta 2. Cell, 126, 543–558.

Barraud, P., Banerjee, S., Mohamed, W.I., Jantsch, M.F. and Allain, F.H.T. (2014) A bimodular nuclear localization signal assembled via an extended double-stranded RNA-binding domain acts as an RNA-sensing signal for transportin 1. Proceedings of the National Academy of Sciences, 111, pE1852–E1861.

Xu, D., Farmer, A. and Chook, Y.M. (2010) Recognition of nuclear targeting signals by Karyopherin-β proteins. Current Opinion in Structural Biology, 20, 782–790.

Kimura, M., Kose, S., Okumura, N., Imai, K., Furuta, M., Sakiyama, N., Tomii, K., Horton, P., Takao, T. and Imamoto, N. (2013) Identification of Cargo Proteins Specific for the Nucleocytoplasmic Transport Carrier Transportin by Combination of an in Vitro Transport System and Stable Isotope Labeling by Amino Acids in Cell Culture (SILAC)-based Quantitative Proteomics. Molecular & Cellular Proteomics: MCP, 12, 145–157.

Twyffels, L., Gueydan, C. and Kruys, V. (2014) Transportin-1 and Transportin-2: Protein nuclear import and beyond. FEBS Letters, 588, 1857–1868.

Carpenter, A.E., Jones, T.R., Lamprecht, M.R., Clarke, C., Kang, I.H., Friman, O., Guertin, D.A., Chang, J.H., Lindquist, R.A., Moffat, J., et al. (2006) CellProfiler: image analysis software for identifying and quantifying cell phenotypes. Genome Biology, 7, R100.

Morrish, T.A., Garcia-Perez, J.L., Stamato, T.D., Taccioli, G.E., Sekiguchi, J. and Moran, J.V. (2007) Endonuclease-independent LINE-1 retrotransposition at mammalian telomeres. Nature, 446, 208–212.

Hunter, SE., Finnegan, EF., Zisoulis, DG., Lovci, MT., Melnik-Martinez, KV., Yeo, GW., Pasquinelli, AE. (2013) Functional genomic analysis of the let-7 regulatory network in Caenorhabditis elegans. PLoS Genet. 9(3):e1003353.

Hogan, D.J., Vincent, T.M., Fish, S., Marcusson, E.G., Bhat, B., Chau, B.N. and Zisoulis, D.G. (2014) Anti-miRs Competitively Inhibit microRNAs in Argonaute Complexes. PLoS ONE, 9, e100951.

Wei, W., Gilbert, N., Ooi, S.L., Lawler, J.F., Ostertag, E.M., Kazazian, H.H., Boeke, J.D. and Moran, J.V. (2001) Human L1 retrotransposition: cis preference versus trans complementation. Mol Cell Biol, 21, 1429–1439.

Agarwal, V., Bell, G.W., Nam, J.-W., Bartel, D.P. and Izaurralde, E. (2015) Predicting effective microRNA target sites in mammalian mRNAs. eLife, 4, e05005.

Cansizoglu, A.E., Lee, B.J., Zhang, Z.C., Fontoura, B.M.A. and Chook, Y.M. (2007) Structure-based design of a pathway-specific nuclear import inhibitor. Nat Struct Mol Biol, 14, 452–454.

Imasaki, T., Shimizu, T., Hashimoto, H., Hidaka, Y., Kose, S., Imamoto, N., Yamada, M. and Sato, M. (2007) Structural Basis for Substrate Recognition and Dissociation by Human Transportin 1. Molecular Cell, 28, 57–67.

Pollard, V.W., Michael, W.M., Nakielny, S., Siomi, M.C., Wang, F. and Dreyfuss, G. (1996) A novel receptor-mediated nuclear protein import pathway. Cell, 86, 985–994.

Nakielny, S., Siomi, M.C., Siomi, H., Michael, W.M., Pollard, V. and Dreyfuss, G. (1996) Transportin: nuclear transport receptor of a novel nuclear protein import pathway. Exp. Cell Res., 229, 261–266.

Fridell, R.A., Truant, R., Thorne, L., Benson, R.E. and Cullen, B.R. (1997) Nuclear import of hnRNP A1 is mediated by a novel cellular cofactor related to karyopherin-beta. J. Cell. Sci., 110 (Pt 11), 1325–1331.

Bonifaci, N., Moroianu, J., Radu, A. and Blobel, G. (1997) Karyopherin beta2 mediates nuclear import of a mRNA binding protein. Proc Natl Acad Sci U S A, 94, 5055–5060.

Lee, B.L. Cansizoglu A.E., Süel, K.E., Louis, T.H., Zhang, Z.C., Chook, Y.M.,(2006) Rules for nuclear localization sequence recognition by Karyopherin beta 2, Cell, 126:543–558

Christ, F., Thys, W., De Rijck, J., Gijsbers, R., Albanese, A., Arosio, D., Emiliani, S., Rain, J.-C., Benarous, R., Cereseto, A., et al. (2008) Transportin-SR2 Imports HIV into the Nucleus. Current Biology, 18, 1192–1202.

Krishnan, L., Matreyek, K.A., Oztop, I., Lee, K., Tipper, C.H., Li, X., Dar, M.J., KewalRamani, V.N. and Engelman, A. (2010) The requirement for cellular transportin 3 (TNPO3 or TRN-SR2) during infection maps to human immunodeficiency virus type 1 capsid and not integrase. J. Virol., 84, 397–406.

Maertens, G.N., Cook, N.J., Wang, W., Hare, S., Gupta, S.S., Oztop, I., Lee, K., Pye, V.E., Cosnefroy, O., Snijders, A.P., et al. (2014) Structural basis for nuclear import of splicing factors by human Transportin 3. Proceedings of the National Academy of Sciences, 111, 2728–2733.

De Iaco, A. and Luban, J. (2011) Inhibition of HIV-1 infection by TNPO3 depletion is determined by capsid and detectable after viral cDNA enters the nucleus. Retrovirology, 8, 1.

De Iaco, A., Santoni, F., Vannier, A., Guipponi, M., Antonarakis, S. and Luban, J. (2013) TNPO3 protects HIV-1 replication from CPSF6-mediated capsid stabilization in the host cell cytoplasm. Retrovirology, 10, 20.

Valle-Casuso, J.C., Di Nunzio, F., Yang, Y., Reszka, N., Lienlaf, M., Arhel, N., Perez, P., Brass, A.L. and Diaz-Griffero, F. (2012) TNPO3 Is Required for HIV-1 Replication after Nuclear Import but prior to Integration and Binds the HIV-1 Core. J. Virol, 86, 5931–5936.

Lee, K., Ambrose, Z., Martin, T.D., Oztop, I., Mulky, A., Julias, J.G., Vandegraaff, N., Baumann, J.G., Wang, R., Yuen, W., et al. (2010) Flexible Use of Nuclear Import Pathways by HIV-1. Cell Host & Microbe, 7, 221–233.

Neumann, M., Bentmann, E., Dormann, D., Jawaid, A., DeJesus-Hernandez, M., Ansorge, O., Roeber, S., Kretzschmar, H.A., Munoz, D.G., Kusaka, H., et al. (2011) FET proteins TAF15 and EWS are selective markers that distinguish FTLD with FUS pathology from amyotrophic lateral sclerosis with FUS mutations. Brain, 134, 2595–2609.

Goodier, J.L., Cheung, L.E. and Kazazian, H.H. (2013) Mapping the LINE1 ORF1 protein interactome reveals associated inhibitors of human retrotransposition. Nucleic Acids Research, 41, 7401–7419.

Pedersen, I.M., Cheng, G., Wieland, S., Volinia, S., Croce, C.M., Chisari, F.V. and David, M. (2007) Interferon modulation of cellular microRNAs as an antiviral mechanism. Nature, 449, 919–922.

Nathans, R., Chu, C.-Y., Serquina, A.K., Lu, C.-C., Cao, H. and Rana, T.M. (2009) Cellular MicroRNA and P Bodies Modulate Host-HIV-1 Interactions. Molecular Cell, 34, 696–709.

Goodier, J.L., Zhang, L., Vetter, M.R. and Kazazian, H.H. (2007) LINE-1 ORF1 Protein Localizes in Stress Granules with Other RNA-Binding Proteins, Including Components of RNA Interference RNA-Induced Silencing Complex. Mol Cell Biol, 27, 6469–6483.

Siomi, M.C., Eder, P.S., Kataoka, N., Wan, L., Liu, Q. and Dreyfuss, G. (1997) Transportin-mediated nuclear import of heterogeneous nuclear RNP proteins. J Cell Biol, 138, 1181–1192.

Rebane, A., Aab, A. and Steitz, J.A. (2004) Transportins 1 and 2 are redundant nuclear import factors for hnRNP A1 and HuR. RNA, 10, 590–599.

Guttinger, S., Muhlhausser, P., Koller-Eichhorn, R., Brennecke, J. and Kutay, U. (2004) From The Cover: Transportin2 functions as importin and mediates nuclear import of HuR. Proc Natl Acad Sci U S A, 101, 2918–2923.

Shi, Z.-M., Wang, J., Yan, Z., You, Y.-P., Li, C.-Y., Qian, X., Yin, Y., Zhao, P., Wang, Y.-Y., Wang, X.-F., et al. (2012) MiR-128 Inhibits Tumor Growth and Angiogenesis by Targeting p70S6K1. PLoS ONE, 7, e32709.

Zhang, Y., Chao, T., Li, R., Liu, W., Chen, Y., Yan, X., Gong, Y., Yin, B., Liu, W., Qiang, B., et al. (2008) MicroRNA-128 inhibits glioma cells proliferation by targeting transcription factor E2F3a.-PubMed-NCBI. J Mol Med, 87, 43–51.

Masri, S., Liu, Z., Phung, S., Wang, E., Yuan, Y.-C. and Chen, S. (2010) The role of microRNA-128a in regulating TGFbeta signaling in letrozole-resistant breast cancer cells.-PubMed-NCBI. Breast Cancer Res Treat, 124, 89–99.

Qian, P., Banerjee, A., Wu, Z.S., Zhang, X., Wang, H., Pandey, V., Zhang, W.J., Lv, X.F., Tan, S., Lobie, P.E., et al. (2012) Loss of SNAIL Regulated miR-128-2 on Chromosome 3p22.3 Targets Multiple Stem Cell Factors to Promote Transformation of Mammary Epithelial Cells. Cancer Res, 72, 6036–6050.

Jin, M., Zhang, T., Liu, C., Badeaux, M.A., Liu, B., Liu, R., Jeter, C., Chen, X., Vlassov, A.V. and Tang, D.G. (2014) miRNA-128 suppresses prostate cancer by inhibiting BMI-1 to inhibit tumor-initiating cells.-PubMed-NCBI. Cancer Res, 74, 4183–4195.

Liu, J., Cao, L., Chen, J., Song, S., Lee, I.H., Quijano, C., Liu, H., Keyvanfar, K., Chen, H., Cao, L.-Y., et al. (2009) Bmi1 regulates mitochondrial function and the DNA damage response pathway. Nature, 459, 387–392.

Sempere, L.F., Freemantle, S., Pitha-Rowe, I., Moss, E., Dmitrovsky, E. and Ambros, V. (2004) Expression profiling of mammalian microRNAs uncovers a subset of brain-expressed microRNAs with possible roles in murine and human neuronal differentiation. Genome Biol, 5, 1.

Fineberg, S.K., Kosik, K.S. and Davidson, B.L. (2009) MicroRNAs potentiate neural development.-PubMed-NCBI. Neuron, 64, 303–309.

Singer, T., McConnell, M.J., Marchetto, M.C.N., Coufal, N.G. and Gage, F.H. (2010) LINE-1 retrotransposons: mediators of somatic variation in neuronal genomes? - PubMed - NCBI. Trends in Neurosciences, 33, 345–354.

Erwin, J.A., Marchetto, M.C. and Gage, F.H. (2014) Mobile DNA elements in the generation of diversity and complexity in the brain. Nat Rev Neurosci, 15, 497–506.

